# Recent evolution of a TET-controlled and DPPA3/STELLA-driven pathway of passive demethylation in mammals

**DOI:** 10.1101/321604

**Authors:** Christopher B. Mulholland, Atsuya Nishiyama, Joel Ryan, Ryohei Nakamura, Merve Yiğit, Ivo M. Glück, Carina Trummer, Weihua Qin, Michael D. Bartoschek, Franziska R. Traube, Edris Parsa, Enes Ugur, Miha Modic, Aishwarya Acharya, Paul Stolz, Christoph Ziegenhain, Michael Wierer, Wolfgang Enard, Thomas Carell, Don C. Lamb, Hiroyuki Takeda, Makoto Nakanashi, Sebastian Bultmann, Heinrich Leonhardt

**Author notes:** Correspondence and requests for material should be addressed to S.B. or H.L. These authors contributed equally (Atsuya Nishiyama, Joel Ryan). These authors jointly supervised this work (Sebastian Bultmann, Heinrich Leonhardt).

## Abstract

Genome-wide DNA demethylation is a unique feature of mammalian development and naïve pluripotent stem cells. So far, it was unclear how mammals specifically achieve global DNA hypomethylation, given the high conservation of the DNA (de-)methylation machinery among vertebrates. We found that DNA demethylation requires TET activity but mostly occurs at sites where TET proteins are not bound suggesting a rather indirect mechanism. Among the few specific genes bound and activated by TET proteins was the naïve pluripotency and germline marker *Dppa3* (*Pgc7, Stella*), which undergoes TDG dependent demethylation. The requirement of TET proteins for genome-wide DNA demethylation could be bypassed by ectopic expression of *Dppa3*. We show that DPPA3 binds and displaces UHRF1 from chromatin and thereby prevents the recruitment and activation of the maintenance DNA methyltransferase DNMT1. We demonstrate that DPPA3 alone can drive global DNA demethylation when transferred to amphibians (*Xenopus*) and fish (medaka), both species that naturally do not have a *Dppa3* gene and exhibit no post-fertilization DNA demethylation. Our results show that TET proteins are responsible for active and - indirectly also for - passive DNA demethylation; while TET proteins initiate local and gene-specific demethylation in vertebrates, the recent emergence of DPPA3 introduced a unique means of genome-wide passive demethylation in mammals and contributed to the evolution of epigenetic regulation during early mammalian development.

## INTRODUCTION

During early embryonic development the epigenome undergoes massive changes. Upon fertilization the genomes of highly specialized cell types — sperm and oocyte — need to be reprogrammed in order to obtain totipotency. This process entails decompaction of the highly condensed gametic genomes and global resetting of chromatin states to confer the necessary epigenetic plasticity required for the development of a new organism^1^. At the same time the genome needs to be protected from the activation of transposable elements (TEs) abundantly present in vertebrate genomes^2^. Activation and subsequent transposition of TEs results in mutations that can have deleterious effects and are passed onto offspring if they occur in the germline during early development^2,3^. The defense against these genomic parasites has shaped genomes substantially^4,5^.

DNA methylation in vertebrates refers to the addition of a methyl group at the C5 position of cytosine to form 5-methylcytosine (5mC). Besides its important role in gene regulation, the most basic function of DNA methylation is the repression and stabilization of TEs and other repetitive sequences^6^. Accordingly, the majority of genomic 5mC is located within these highly abundant repetitive elements. Global DNA methylation loss triggers the derepression of transposable and repetitive elements, which leads to genomic instability and cell death, highlighting the crucial function of vertebrate DNA methylation^7–12^. Hence, to ensure constant protection against TE reactivation, global DNA methylation levels remain constant throughout the lifetime of non-mammalian vertebrates^13–16^. Paradoxically, mammals specifically erase DNA methylation during preimplantation development^17,18^, a process that would seemingly expose the developing organism to the risk of genomic instability through the activation of TEs. DNA methylation also acts as an epigenetic barrier to restrict and stabilize cell fate decisions and thus constitutes a form of epigenetic memory. The establishment of pluripotency in mammals requires the erasure of epigenetic memory and as such, global hypomethylation is a defining characteristic of pluripotent cell types including naïve embryonic stem cells (ESCs), primordial germ cells (PGCs), and induced pluripotent stem cells (iPSCs)^19^.

In vertebrates, DNA methylation can be reversed to unmodified cytosine by two mechanisms; either actively by Ten-eleven translocation (TET) dioxygenase-mediated oxidation of 5mC in concert with the base excision repair machinery^20–23^ or passively by a lack of functional DNA methylation maintenance during the DNA replication cycle^24,25^. In mammals, both active and passive demethylation pathways have been suggested to be involved in the establishment of global demethylation during preimplantation development^26–32^. Intriguingly however, whereas hypomethylated states during development are mammal-specific, TET-proteins as well as the DNA methylation machinery are highly conserved among all vertebrates^33,34^. This discrepancy implies the existence of so far unknown mammalian-specific pathways and factors controlling the establishment and maintenance of genomic hypomethylation.

Here, we use mouse embryonic stem cells (ESCs) cultured in conditions promoting naïve pluripotency^35–37^ as a model to study global DNA demethylation in mammals. By dissecting the contribution of the catalytic activity of TET1 and TET2 we show that TET-mediated active demethylation drives the expression of the Developmental pluripotency-associated protein 3 (DPPA3/PGC7/STELLA). We show that DPPA3 directly binds UHRF1 prompting its release from chromatin, resulting in the inhibition of maintenance methylation and global passive demethylation. Although only found in mammals, DPPA3 can also induce global demethylation in non-mammalian vertebrates. In summary, we described a novel TET-controlled and DPPA3-driven pathway for passive demethylation in naïve pluripotency in mammals.

## RESULTS

### TET1 and TET2 indirectly protect the naïve genome from hypermethylation

Mammalian TET proteins, TET1, TET2, and TET3, share a conserved catalytic domain and the ability to oxidize 5mC but exhibit distinct expression profiles during development^38^. Naïve ESCs and the inner cell mass (ICM) of the blastocyst from which they are derived feature high expression of *Tet1* and *Tet2* but not *Tet3*^27,39–41^. To dissect the precise contribution of TET-mediated active DNA demethylation to global DNA hypomethylation in naïve pluripotency we generated isogenic *Tet1* (T1CM) and *Tet2* (T2CM) single as well as *Tet1/Tet2* (T12CM) double catalytic mutant mouse ESC lines using CRISPR/Cas-assisted gene editing (Supplementary Fig. 1). We confirmed the inactivation of TET1 and TET2 activity by measuring the levels of 5-hydroxymethylcytosine (5hmC), the product of TET-mediated oxidation of 5mC^20^ (Supplementary Fig. 1i). While the loss of either *Tet1* or *Tet2* catalytic activity significantly reduced 5hmC levels, inactivation of both TET1 and TET2 resulted in the near total loss of 5hmC in naïve ESCs (Supplementary Fig. 1i) indicating that TET1 and TET2 account for the overwhelming majority of cytosine oxidation in naïve ESCs. We then used reduced representation bisulfite sequencing (RRBS) to determine the DNA methylation state of T1CM, T2CM, and T12CM ESCs as well as wild-type (wt) ESCs. All *Tet* catalytic mutant (T1CM, T2CM and T1CM) cell lines exhibited severe DNA hypermethylation throughout the genome including promoters, gene bodies, and repetitive elements (Fig. 1a,b). The increase in DNA methylation was particularly pronounced at LINE-1 (L1) elements of which 97%, 98%, and 99% were significantly hypermethylated in T1CM, T2CM, and T12CM ESCs, respectively (Supplementary Fig. 2a). This widespread DNA hypermethylation was reminiscent of the global increase in DNA methylation accompanying the transition of naïve ESCs to primed epiblast-like cells (EpiLCs)^40,42,43^, which prompted us to investigate whether the hypermethylation present in T1CM, T2CM, and T12CM ESCs represents a premature acquisition of a more differentiated DNA methylation signature. In line with this hypothesis, *Tet* catalytic mutant ESCs displayed DNA methylation levels similar or higher than those of wt EpiLCs (Supplementary Fig. 2b). Moreover, hierarchical clustering and principal component analyses (PCA) of the RRBS data revealed that ESCs from *Tet* catalytic mutants clustered closer to wt EpiLCs than wt ESCs (Fig. 1c and Supplementary Fig. 2c). In fact, the vast majority of significantly hypermethylated CpGs in *Tet* catalytic mutant ESCs overlapped with those normally gaining DNA methylation during the exit from naïve pluripotency (Fig. 1d). However, T1CM, T2CM, and T12CM transcriptomes clearly clustered by differentiation stage indicating that the acquisition of an EpiLC-like methylome was not due to premature differentiation (Supplementary Fig. 2d). Intriguingly, we found that the majority of sites hypermethylated in *Tet* catalytic mutant ESCs are not bound by TET1 or TET2 (Fig. 1e,f) suggesting that the catalytic activity of TET1 and TET2 maintains the hypomethylated state of the naïve methylome by indirect means.

**Fig. 1:**
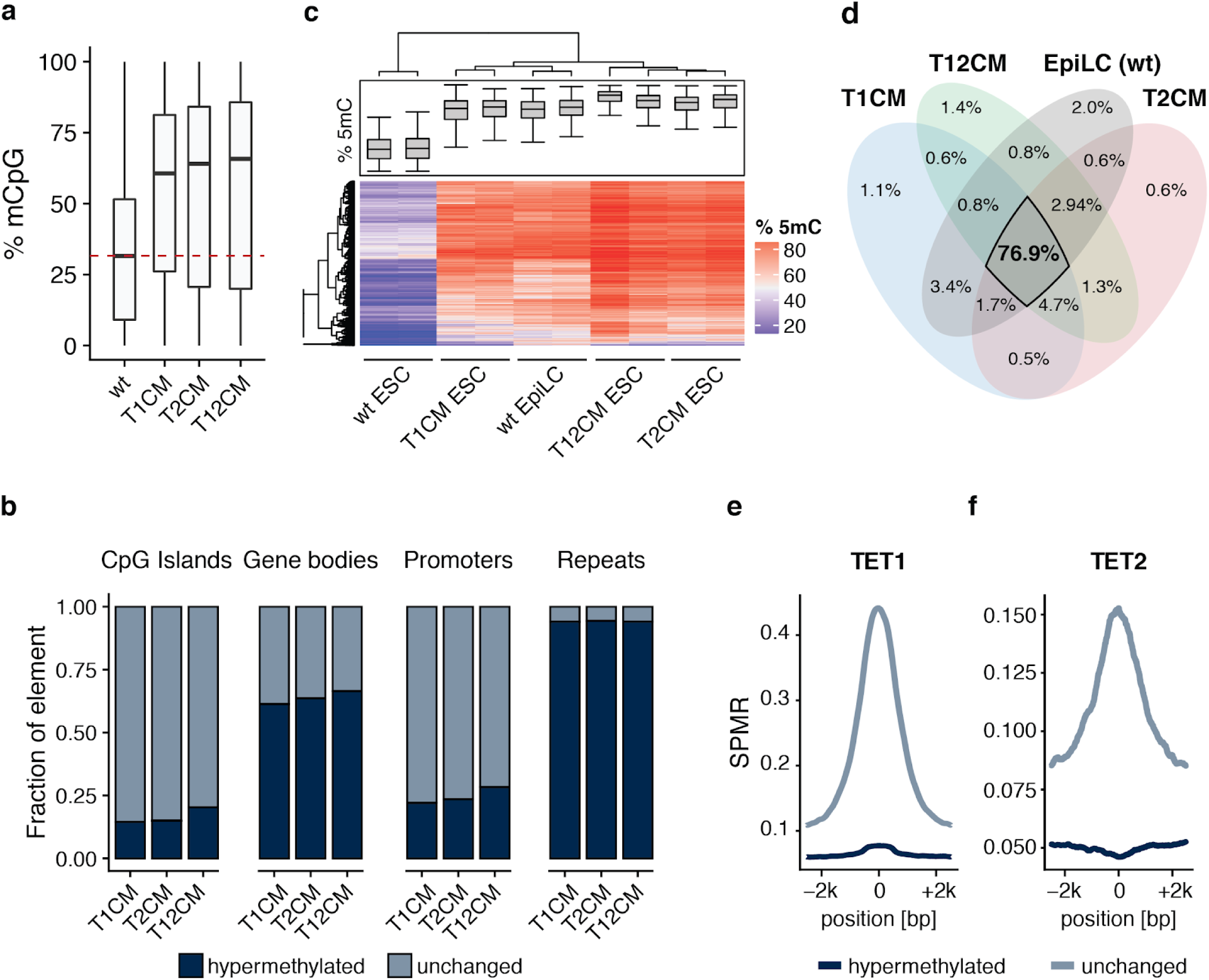
TET1 and TET2 prevent hypermethylation of the naïve genome. **a**, Loss of TET catalytic activity leads to global DNA hypermethylation. Percentage of total 5mC as measured by RRBS. **b**, Loss of TET catalytic activity leads to widespread DNA hypermethylation especially at repetitive elements. Relative proportion of DNA hypermethylation (q value < 0.05; absolute methylation difference > 20%) at each genomic element in T1CM, T2CM, and T12CM ESCs compared to wt ESCs. **c**, Heat map of the hierarchical clustering of the RRBS data depicting the top 2000 most variable 1kb tiles during differentiation of wt ESCs to EpiLCs. **d**, Venn diagram depicting the overlap of hypermethylated (compared to wt ESCs; q value < 0.05; absolute methylation difference > 20%) sites among T1CM, T2CM, and T12CM ESCs and wt EpiLCs. **e,f**, TET binding is not associated with DNA hypermethylation in TET mutant ESCs. Occupancy of (**e**) TET1^44^ and (**f**) TET2^45^ over 1 kb tiles hypermethylated (dark blue) or unchanged (light blue) in T1CM and T2CM ESCs, respectively. In the boxplots in **a-c**, horizontal black lines within boxes represent median values, boxes indicate the upper and lower quartiles, and whiskers indicate the 1.5 interquartile range.

### TET1 and TET2 control *Dppa3* expression in a catalytically dependent manner

To understand how TET1 and TET2 indirectly promote demethylation of the naïve genome, we examined the expression of the enzymes involved in DNA methylation. Loss of TET catalytic activity was not associated with changes in the expression of *Dnmt1, Uhrf1, Dnmt3a*, and *Dnmt3b*, indicating the hypermethylation in *Tet* catalytic mutant ESCs is not a result of the DNA methylation machinery being upregulated (Fig. 2a). To identify candidate factors involved in promoting global hypomethylation, we compared the transcriptome of hypomethylated wild-type ESCs with those of hypermethylated cells, which included wt EpiLCs as well as T1CM, T2CM, and T12CM ESCs (Fig. 2b). Among the 14 genes differentially expressed in hypermethylated cell lines, the naïve pluripotency factor, *Dppa3* (also known as *Stella* and *Pgc7*), was an interesting candidate due to its reported involvement in the regulation of global DNA methylation in germ cell development and oocyte maturation^46–49^. In addition, *Dppa3* is also a direct target of PRDM14, a PR domain-containing transcriptional regulator known to promote the DNA hypomethylation associated with naïve pluripotency^36,50–52^ (Fig. 2e). While normally highly expressed in naïve ESCs and only downregulated upon differentiation^53,54^, *Dppa3* was prematurely repressed in T1CM, T2CM, and T12CM ESCs (Fig. 2d). The significantly reduced expression of *Dppa3* in TET mutant ESCs was accompanied by significant hypermethylation of the *Dppa3* promoter (Fig. 2e), consistent with reports demonstrating *Dppa3* to be one of the few pluripotency factors regulated by promoter methylation *in vitro* and *in vivo*^37,53–55^. In contrast to the majority of genomic sites gaining methylation in TET mutant ESCs (Fig. 1e,f), hypermethylation at the *Dppa3* locus occurred at sites bound by both TET1 and TET2 (Fig. 2e)^44,45^. This hypermethylation overlapped with regions at which the TET oxidation product 5-carboxylcytosine (5caC) accumulates in Thymine DNA glycosylase (TDG)-knockdown ESCs (Fig. 2e)^56^, indicating that the Dppa3 locus is a direct target of TET/TDG-mediated active DNA demethylation in ESCs.

**Fig. 2:**
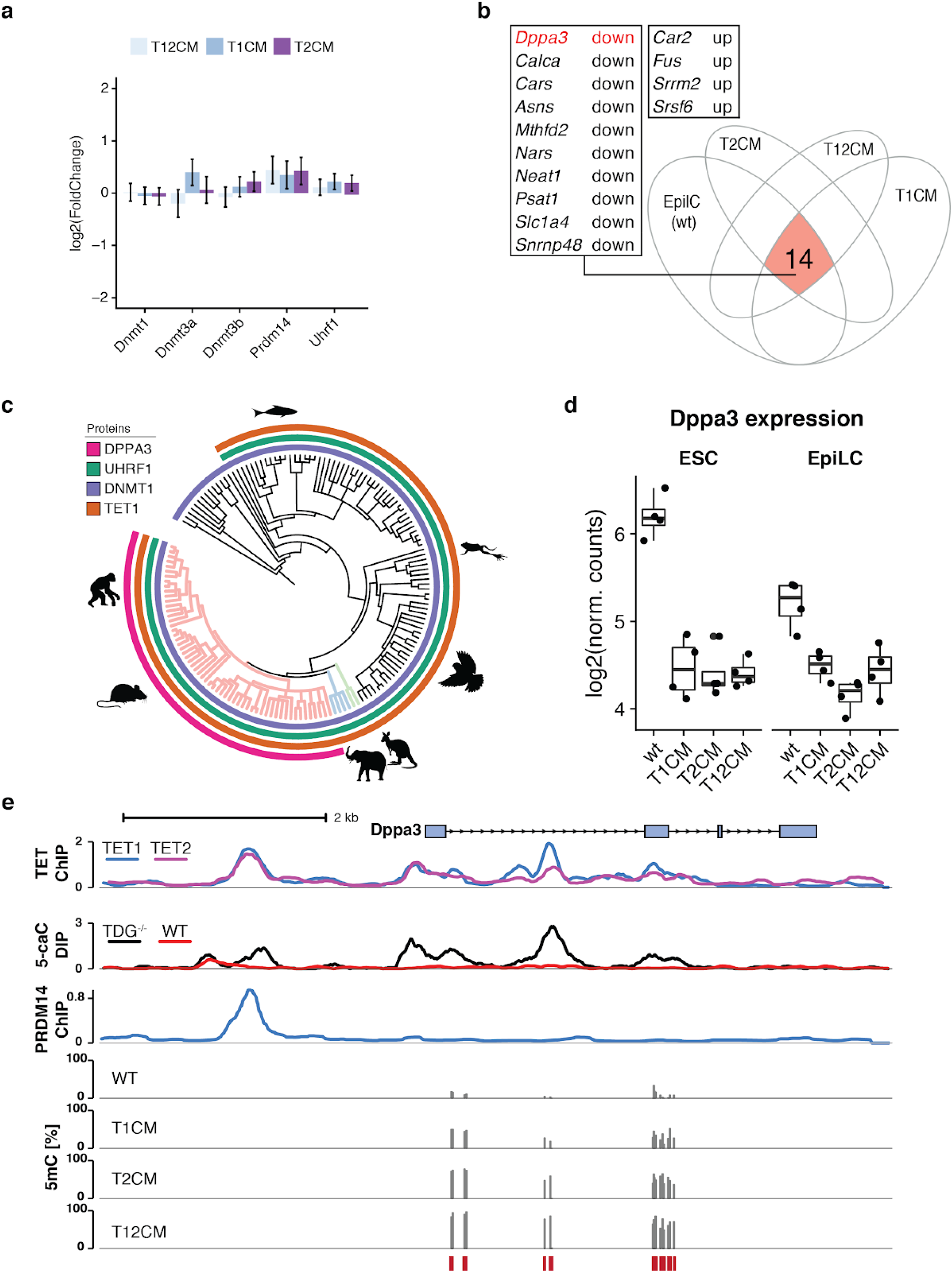
TET1 and TET2 catalytic activity is necessary for *Dppa3* expression. **a**, Expression of genes involved in regulating DNA methylation levels in T1CM, T2CM, and T12CM ESCs as assessed by RNA-seq. Expression is given as the log_2_ fold-change compared to wt ESCs. Error bars indicate mean ± SD, *n* = 4 biological replicates. No significant changes observable (Likelihood ratio test). **b**, *Dppa3* is downregulated upon loss of TET activity and during differentiation. Venn diagram depicting the overlap (red) of genes differentially expressed (compared to wt ESCs; adjusted p < 0.05) in T1CM, T2CM, T12CM ESCs, and wt EpiLCs. **c**, Phylogenetic tree of TET1, DNMT1, UHRF1, and DPPA3 in metazoa. **d**, *Dppa3* expression levels as determined by RNA-seq in the indicated ESC and EpiLC lines (*n* = 4 biological replicates). **e**, TET proteins bind and actively demethylate the *Dppa3* locus. Genome browser view of the *Dppa3* locus with tracks of the occupancy (Signal pileup per million reads; (SPMR)) of TET1^44^, TET2^45^, and PRDM14^51^ in wt ESCs, 5caC enrichment in wt vs. TDG^-/-^ ESCs^56^, and 5mC (%) levels in wt, T1CM, T2CM, and T12CM ESCs (RRBS). Red bars indicate CpGs covered by RRBS. In **d**, boxplots horizontal black lines within boxes represent median values, boxes indicate the upper and lower quartiles, and whiskers indicate the 1.5 interquartile range. P-values were calculated using Welch’s two-sided t-test comparing *Tet* catalytic mutants to their corresponding wt: ** P < 0.01; *** P < 0.001.

PRDM14 has been shown to recruit TET1 and TET2 to sites of active demethylation and establish global hypomethylation in naïve pluripotency^36,40,51,52,57^. As the expression of *Prdm14* was not altered in *Tet* catalytic mutant ESCs (Fig. 2a), we analysed PRDM14 occupancy at the *Dppa3* locus using publicly available ChIP-seq data^51^. This analysis revealed that PRDM14 binds the same upstream region of *Dppa3* occupied by TET1 and TET2 (Fig. 2e). Taken together, these data suggest that TET1 and TET2 are recruited by PRDM14 to maintain the expression of *Dppa3* by active DNA demethylation. Strikingly, systematic analysis of public databases revealed that while the DNA (de)methylation machinery (DNMTs, UHRF1, TETs) is conserved among vertebrates, *Dppa3* is only present in mammals potentially representing a novel pathway that regulates mammalian-specific global hypomethylation in naïve pluripotency (Fig. 2c).

### DPPA3 acts downstream of TET1 and TET2 and is required to safeguard the naïve methylome

DPPA3 has been reported to both prevent or promote DNA demethylation depending on the developmental time points^46,48,49,58–62^. However, the function of DPPA3 in the context of naïve pluripotency, for which it is a well-established marker gene^53^, has yet to be explored. We first sought to characterize the relationship between the high expression of *Dppa3* and DNA hypomethylation both accompanying naïve pluripotency. To this end, we established isogenic *Dppa3* knockout (Dppa3KO) mouse ESCs using CRISPR/Cas (Supplementary Fig. 3a-c) and profiled their methylome by RRBS. Loss of DPPA3 led to severe global hypermethylation (Fig. 3a) with substantial increases in DNA methylation observed across all genomic elements (Supplementary Fig. 3d). Repetitive sequences and TEs, in particular, were severely hypermethylated including 98% of L1 elements (Supplementary Fig. 3d). A principal component analysis of the RRBS data revealed that Dppa3KO ESCs clustered closer to wt EpiLCs and *Tet* catalytic mutant ESCs rather than wt ESCs (Fig. 3b). Furthermore, we observed a striking overlap of hypermethylated CpGs between *Tet* catalytic mutant and Dppa3KO ESCs (Fig. 3c), suggesting that DPPA3 and TETs promote demethylation at largely the same targets. A closer examination of the genomic distribution of overlapping hypermethylation in *Tet* catalytic mutant and Dppa3KO ESCs revealed that the majority (∼90%) of common targets reside within repetitive elements (Fig. 3d, Supplementary Fig. 3e,f) and are globally correlated with heterochromatic histone modifications (Supplementary Fig. 3h). In contrast, only half of the observed promoter hypermethylation among all cell lines was dependent on DPPA3 (classified as “common”, Fig. 3d and Supplementary Fig. 3e-g). This allowed us to identify a set of strictly TET-dependent promoters (N = 1573) (Fig. 3d, Supplementary Fig. 3f and Supplementary Table 1), which were enriched for developmental genes (Fig 3e and Supplementary Table 2). Intriguingly, these TET-specific promoters contained genes (such as *Pax6, Foxa1* and *Otx2*) that have recently been shown to be conserved targets of TET-mediated demethylation during *Xenopus*, zebrafish and mouse development^63^.

**Fig. 3:**
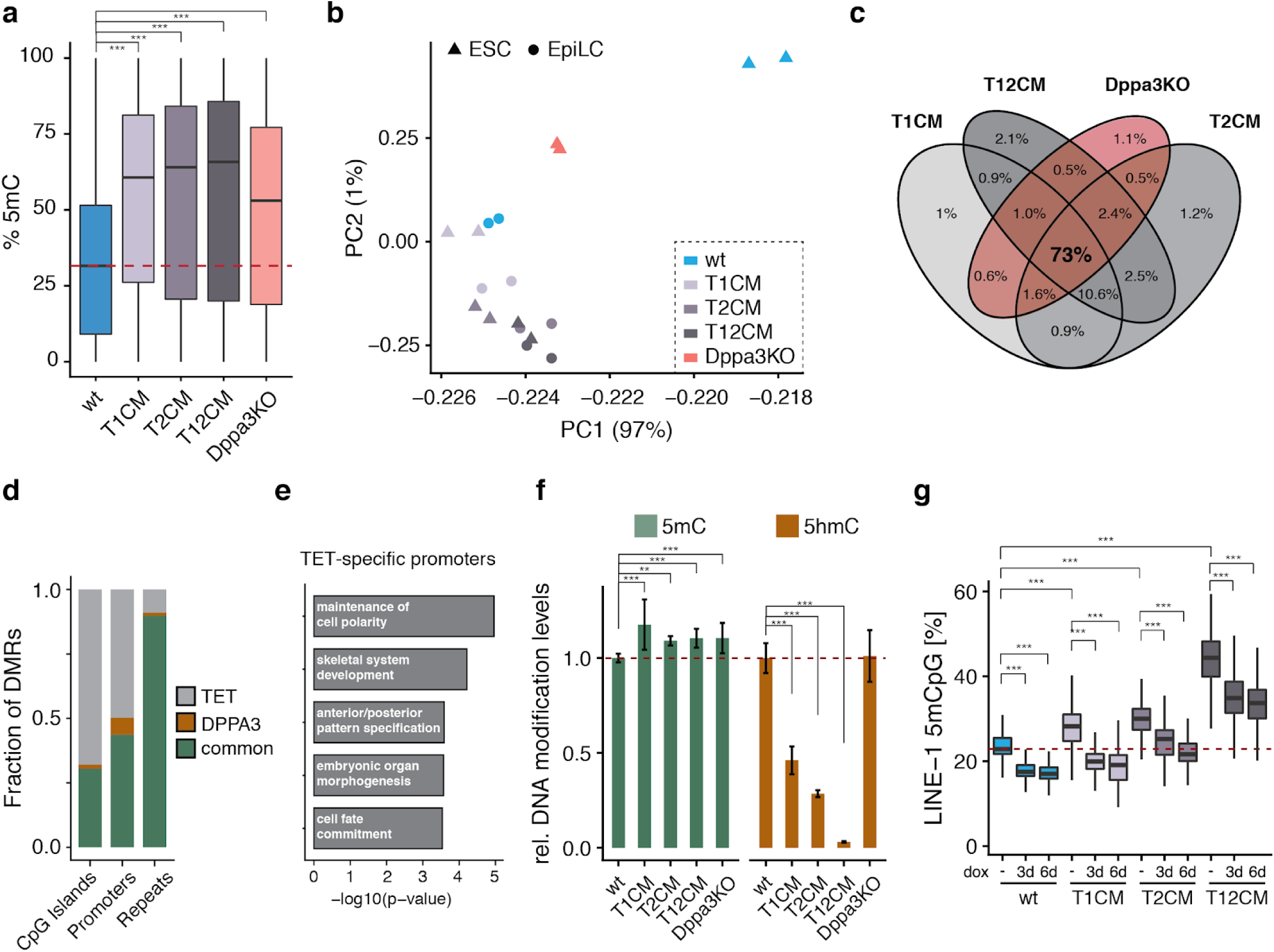
DPPA3 acts downstream of TET1 and TET2 to establish and preserve global hypomethylation. **a**, *Dppa3* loss results in global hypermethylation. Percentage of total 5mC as measured by RRBS. **b**, *Dppa3* prevents the premature acquisition of a primed methylome. Principal component (PC) analysis of RRBS data from wt, T1CM, T2CM and T12CM ESCs, wt EpiLCs and Dppa3KO ESCs. **c**, DPPA3 and TET proteins promote demethylation of largely similar targets. Venn Diagrams depicting the overlap of hypermethylated sites among T1CM, T2CM, T12CM, and Dppa3KO ESCs. **d**, *Dppa3* protects mostly repeats from hypermethylation. Fraction of hypermethylated genomic elements classified as TET-specific (only hypermethylated in TET mutant ESCs), DPPA3-specific (only hypermethylated in Dppa3KO ESCs), or common (hypermethylated in TET mutant and Dppa3KO ESCs). **e**, Gene ontology (GO) terms associated with promoters specifically dependent on TET activity; adjusted p-values calculated using Fisher’s exact test followed by Benjamini-Hochberg correction for multiple testing. **f**, TET activity remains unaffected in Dppa3KO ESCs. Relative DNA modification levels for 5-methylcytosine (5mC) and 5-hydroxymethylcytosine (5hmC) as measured by mass spectrometry (LC-MS/MS). Error bars indicate mean ± SD calculated from *n* > 3 biological replicates. **g**, *Dppa3* expression can rescue the hypermethylation in TET mutant ESCs. DNA methylation levels at LINE-1 elements (%) as measured by bisulfite sequencing 0, 3, or 6 days after doxycycline (dox) induction of *Dppa3* expression. In the boxplots in **a, g**, horizontal black lines within boxes represent median values, boxes indicate the upper and lower quartiles, and whiskers indicate the 1.5 interquartile range. The dashed red line indicates the median methylation level of wt ESCs. In **a, f**, and **g**, P-values were calculated using Welch’s two-sided t-test: ** P < 0.01; *** P < 0.001.

DPPA3 appeared to act downstream of TETs as the global increase in DNA methylation in Dppa3KO ESCs was not associated with a reduction in 5hmC levels nor with a downregulation of TET family members (Fig. 3f and Supplementary Fig. 3i). In support of this notion, inducible overexpression of *Dppa3* (Supplementary Fig. 3j-l) completely rescued the observed hypermethylation phenotype at LINE-1 elements in T1CM as well as T2CM ESCs and resulted in a significant reduction of hypermethylation in T12CM cells (Fig. 3g). Strikingly, prolonged induction of *Dppa3* even resulted in hypomethylation in wild-type as well as T1CM ESCs (Fig. 3g). Collectively, these results show that TET1 and TET2 activity contributes to genomic hypomethylation in naïve pluripotency by direct and indirect pathways, the active demethylation of developmental promoters and the passive, DPPA3-mediated global demethylation.

### TET-dependent expression of DPPA3 regulates UHRF1 subcellular distribution and controls DNA methylation maintenance in embryonic stem cells

To investigate the mechanism underlying the regulation of global DNA methylation patterns by DPPA3 we first generated an endogenous DPPA3-HALO fusion ESC line to monitor the localization of DPPA3 throughout the cell cycle (Supplementary Fig. 4a,c). Recent studies have shown that DPPA3 binds H3K9me2^61^ and that in oocytes its nuclear localization is critical to inhibit the activity of UHRF1^49^, a key factor for maintaining methylation. Expecting a related mechanism to be present in ESCs, we were surprised to find a strong cytoplasmic localization of DPPA3 in ESCs (Fig. 4a). Furthermore, DPPA3 did not bind to mitotic chromosomes indicating a low or absent chromatin association of DPPA3 in ESCs (Fig. 4a). To further understand the mechanistic basis of DPPA3-dependent DNA demethylation in ESCs, we performed FLAG-DPPA3 pulldowns followed by liquid chromatography tandem mass spectrometry (LC MS-MS) to profile the DPPA3 interactome in naïve ESCs. Strikingly, among the 303 significantly enriched DPPA3 interaction partners identified by mass spectrometry, we found UHRF1 and DNMT1 (Fig. 4b and Supplementary Table 3), the core components of the DNA maintenance methylation machinery^64,65^. A reciprocal immunoprecipitation of UHRF1 confirmed its interaction with DPPA3 in ESCs (Supplementary Fig. 4f). Furthermore, GO analysis using the top 131 interactors of DPPA3 showed the two most enriched GO terms to be related to DNA methylation (Supplementary Table 4). These findings were consistent with previous studies implicating DPPA3 in the regulation of maintenance methylation^47,49^. In addition, we also detected multiple members of the nuclear transport machinery indicating that DPPA3 might undergo nuclear shuttling in ESCs (highlighted in purple, Fig. 4b and Supplementary Table 3), which prompted us to investigate whether DPPA3 influences the subcellular localization of UHRF1. Surprisingly, biochemical fractionation experiments revealed UHRF1 to be present in both the nucleus and cytoplasm of naïve wt ESCs (Supplementary Fig. 4e). Despite comparable total UHRF1 protein levels in wt and Dppa3KO ESCs (Supplementary Fig. 4g), loss of DPPA3 completely abolished the cytoplasmic fraction of UHRF1 (Supplementary Fig. 4e).

**Fig. 4:**
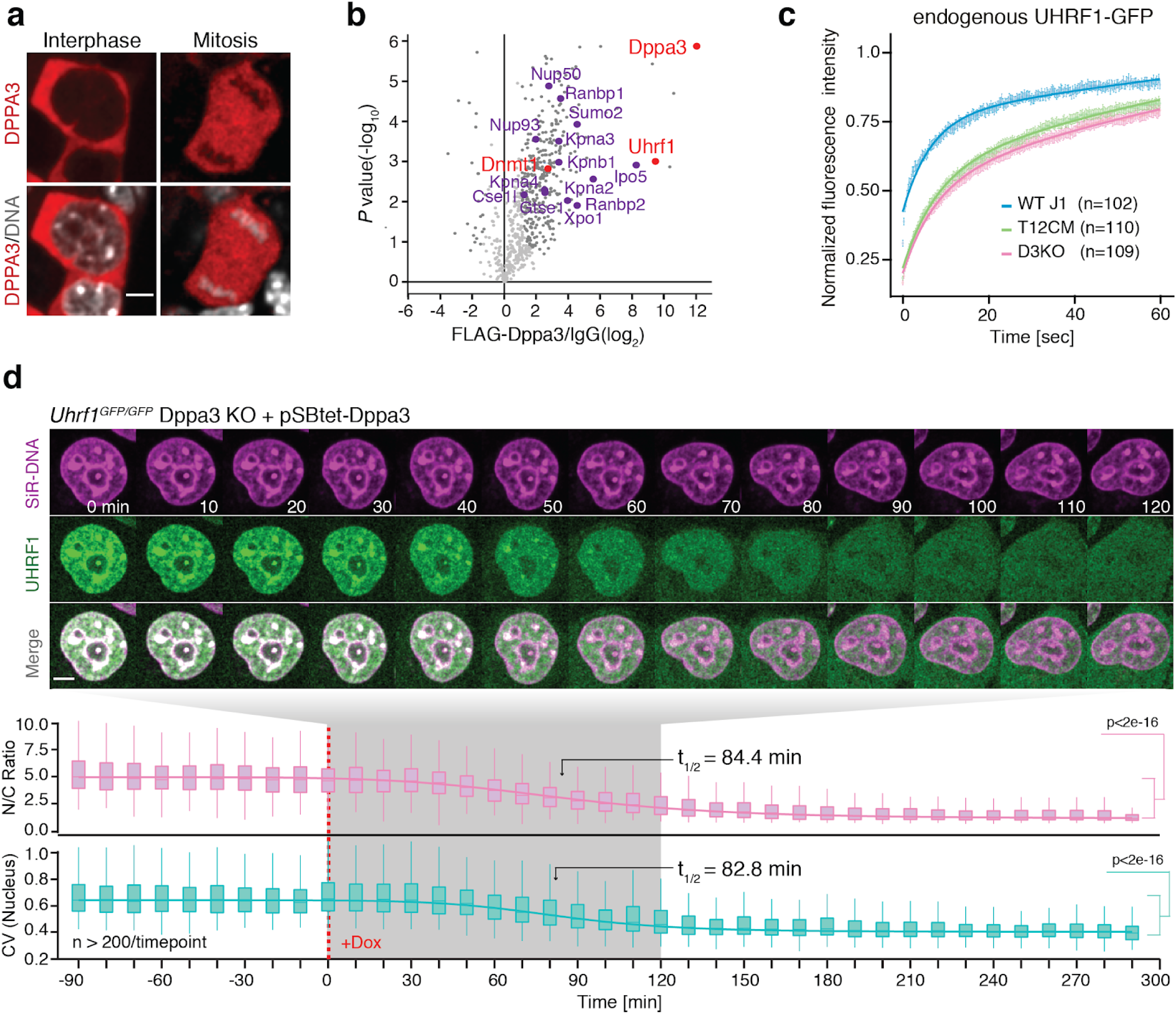
DPPA3 prevents UHRF1 chromatin binding to impede maintenance methylation in embryonic stem cells. **a**, DPPA3 is primarily localized to the cytoplasm of ESCs. Images illustrating the localization of endogenous DPPA3-HALO in live ESCs counterstained with SiR-Hoechst (DNA). Scale bar: 5 μm. **b**, DPPA3 interacts with endogenous UHRF1 in ESCs. Volcano plot from DPPA3-FLAG pulldowns in ESCs. Dark grey dots: significantly enriched proteins (FDR < 0.05). Red dots: proteins involved in DNA methylation regulation. Purple dots: proteins involved in nuclear transport. anti-FLAG antibody: *n* = 3 biological replicates, IgG control antibody: *n* = 3 biological replicates. Statistical significance determined by performing a Student’s t-test with a permutation-based FDR of 0.05 and an additional constant S0 = 1. **c**, Loss of DPPA3 leads to increased UHRF1 chromatin binding. FRAP analysis of endogenous UHRF1-GFP. Each genotype comprises the combined single-cell data from two independent clones acquired in two independent experiments. **d**, Normal UHRF1 localization can be rescued by ectopic *Dppa3* expression. Localization dynamics of endogenous UHRF1-GFP in response to *Dppa3* induction in U1G/D3KO + pSBtet-D3 ESCs with confocal time-lapse imaging over 8 h (10 min intervals). *t = 0* corresponds to start of *Dppa3* induction with doxycycline (+Dox). (*top panel*) Representative images of UHRF1-GFP and DNA (SiR-Hoechst stain) throughout confocal time-lapse imaging. Scale bar: 5 μm. (middle panel) Nucleus to cytoplasm ratio (N/C Ratio) of endogenous UHRF1-GFP signal. (bottom panel) Coefficient of variance (CV) of endogenous UHRF1-GFP intensity in the nucleus. (*middle and bottom panel*) N/C Ratio and CV values: measurements in *n* > 200 single cells per time point, acquired at *n* = 16 separate positions. Curves represent fits of four parameter logistic (4PL) functions to the N/C Ratio (pink line) and CV (green line) data. Live-cell imaging was repeated 3 times with similar results. In **c**, the mean fluorescence intensity of *n* cells (indicated in the plots) at each timepoint are depicted as shaded dots. Error bars indicate mean ± SEM. Curves (solid lines) indicate double-exponential functions fitted to the FRAP data. In the boxplots in **d**, darker horizontal lines within boxes represent median values. The limits of the boxes indicate upper and lower quartiles, and whiskers indicate the 1.5-fold interquartile range. P-values based on Welch’s two-sided t-test.

As maintenance DNA methylation critically depends on the correct targeting and localization of UHRF1 within the nucleus^66–69^, we asked whether TET-dependent regulation of DPPA3 might affect the subnuclear distribution of UHRF1. To this end, we tagged endogenous UHRF1 with GFP in wild-type (U1G/wt) as well as Dppa3KO and T12CM ESCs (U1G/Dppa3KO and U1G/T12CM, respectively) enabling us to monitor UHRF1 localization dynamics in living cells (Supplementary Fig. 4b-d). Whereas UHRF1-GFP localized to both the nucleus and cytoplasm of wt ESCs, UHRF1-GFP localization was solely nuclear in Dppa3KO and T12CM ESCs (Supplementary Fig. 4h,i). In addition, UHRF1 appeared to display a more diffuse localization in wt ESCs compared to Dppa3KO and T12CM ESCs, in which we observed more focal patterning of UHRF1 particularly at heterochromatic foci (Supplementary Fig. 4h). To quantify this observation, we calculated the coefficient of variation (CV) of nuclear UHRF1-GFP among wt, Dppa3KO, and T12CM ESCs. The CV of a fluorescent signal correlates with its distribution, where low CV values reflect more homogenous distributions and high CV values correspond to more heterogeneous distributions^70,71^. Indeed, the pronounced focal accumulation of UHRF1-GFP observed in Dppa3KO and T12CM ESCs corresponded with a highly significant increase in the CV values of nuclear UHRF1-GFP compared with wt ESCs (Supplementary Fig. 4h,i).

To assess whether these differences in nuclear UHRF1 distribution reflected altered chromatin binding, we used fluorescence recovery after photobleaching (FRAP) to study the dynamics of nuclear UHRF1-GFP in wt, Dppa3KO, and T12CM ESCs. Our FRAP analysis revealed markedly increased UHRF1 chromatin binding in both Dppa3KO and T12CM ESCs as evidenced by the significantly slower recovery of UHRF1-GFP in these cell lines compared to wt ESCs (Fig. 4c and Supplementary Fig. 4j,k). Additionally, these data demonstrated increased UHRF1 chromatin binding to underlie the more heterogenous nuclear UHRF1 distributions in Dppa3KO and T12CM ESCs. Interestingly, although strongly reduced compared to wt ESCs, UHRF1 mobility was slightly higher in T12CM ESCs than Dppa3KO ESCs, consistent with a severe but not total loss of DPPA3 in the absence of TET activity (Supplementary Fig. 4l). Induction of *Dppa3* rescued the cytoplasmic fraction of UHRF1 (N/C Ratio: Fig. 4d) as well as the diffuse localization of nuclear UHRF1 in Dppa3KO ESCs (CV: Fig. 4d), which reflected a striking increase in the mobility of residual nuclear UHRF1-GFP as assessed by FRAP (Supplementary Figs. 4m and 5a,b). This analysis also revealed that UHRF1’s nucleocytoplasmic translocation and the inhibition of chromatin binding followed almost identical kinetics (N/C t_1/2_ = 84.4 min; CV t_1/2_ = 82.8) (Fig. 4d). UHRF1 is required for the proper targeting of DNMT1 to DNA replication sites and therefore essential for DNA methylation maintenance^64,72^. We observed a marked reduction of both UHRF1 and DNMT1 at replication foci upon induction of *Dppa3*, indicating that DPPA3 promotes hypomethylation in naïve ESCs by impairing DNA methylation maintenance (Supplementary Fig. 5c,d). Ectopic expression of DPPA3 not only altered the subcellular distribution of endogenous UHRF1 in mouse ESCs (Fig. 4d and Supplementary Fig. 5e) but also in human ESCs suggesting evolutionary conservation of this mechanism among mammals (Supplementary Fig. 5f,g). Collectively our results demonstrate that TET-proteins control both the subcellular localization and chromatin binding of UHRF1 via the regulation of DPPA3 levels in naïve ESCs. Furthermore, these data show that DPPA3 is both necessary and sufficient for ensuring the nucleocytoplasmic translocation, diffuse nuclear localization, and attenuated chromatin binding of UHRF1 in ESCs.

### DPPA3-mediated inhibition of UHRF1 chromatin binding causes hypomethylation and is attenuated by nuclear export

Our results demonstrated cytoplasmic accumulation of UHRF1 and the disruption of its focal nuclear patterning to occur with almost identical kinetics upon induction of *Dppa3* expression (Fig. 4d). In principle, either a decrease in nuclear UHRF1 concentration or the impaired chromatin loading of UHRF1 would on their own be sufficient to impair maintenance DNA methylation^68,73^. To dissect these two modes and their contribution to the inhibition of maintenance methylation in naïve ESCs, we generated inducible *Dppa3*-mScarlet expression cassettes (Supplementary Fig. 6a) harboring mutations to residues described to be critical for its nuclear export (ΔNES)^48^ and the interaction with UHRF1 (KRR and R107E)^49^, as well as truncated forms of DPPA3 found in zygotes, 1-60 and 61-150^62^ (Fig. 5a). After introducing these *Dppa3* expression cassettes into U1GFP/Dppa3KO ESCs, we used live-cell imaging to track each DPPA3 mutant’s localization and ability to rescue the loss of DPPA3 (Fig. 5b). Whereas DPPA3-ΔNES and DPPA3 61-150, both lacking a functional nuclear export signal, were sequestered to the nucleus (Fig. 5b), DPPA3-WT, DPPA3-KRR, DPPA3-R107E, and DPPA3 1-60 mutants localized primarily to the cytoplasm (Fig. 5b) and recapitulated the localization of endogenous DPPA3 in naïve ESCs (Fig. 4a). Regardless, all tested DPPA3 mutants failed to efficiently reestablish nucleocytoplasmic translocation of UHRF1 (Fig. 5b and Supplementary Fig. 6b), indicating that DPPA3 requires both the capacity to interact with UHRF1 as well as a functional nuclear export signal to promote nucleocytoplasmic shuttling of UHRF1 in naïve ESCs.

**Fig. 5:**
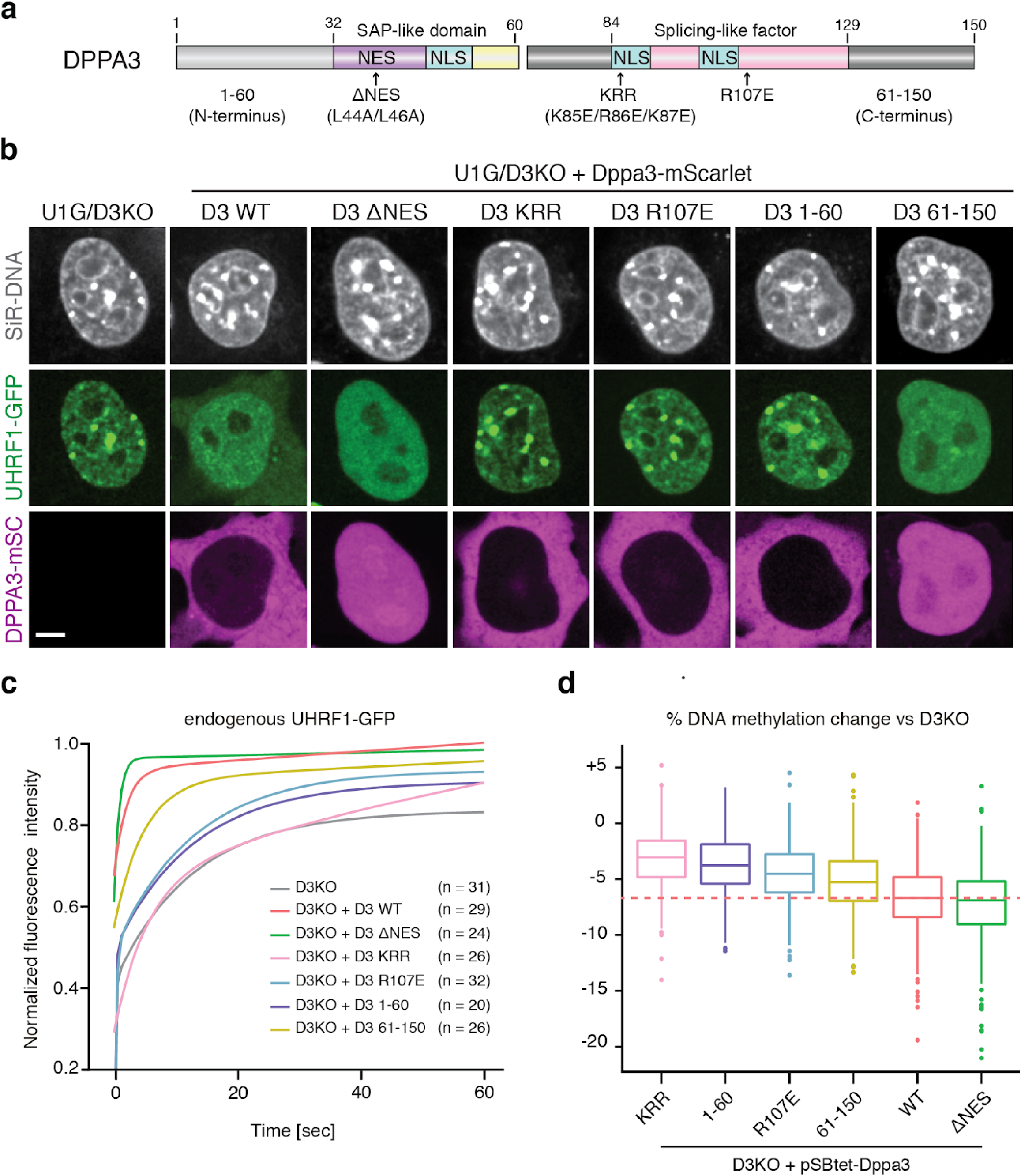
DPPA3 inhibits maintenance DNA methylation by impairing UHRF1 chromatin binding. **a**, Schematic illustration of murine DPPA3 with the nuclear localization signals (NLS), nuclear export signal (NES), and predicted domains (SAP-like and splicing factor-like^74^) annotated. For the DPPA3 mutant forms used in this study, point mutations are indicated with arrows (ΔNES, KRR, R107E) and the two truncations are denoted by the middle break (1-60, left half; 61-150, right half). **b,c**, Nuclear export and the C-terminus of DPPA3 are dispensable for disrupting focal UHRF1 patterning and chromatin binding in ESCs. **b**, Representative confocal images illustrating the localization of endogenous UHRF1-GFP and the indicated mDPPA3-mScarlet fusions in live U1G/D3KO + pSB-D3-mSC ESCs. DNA counterstain: SiR-Hoechst. Scale bar: 5 μm. **c**, FRAP analysis of endogenous UHRF1-GFP in U1G/D3KO ESCs expressing the indicated mutant forms of DPPA3. FRAP Curves (solid lines) indicate double-exponential functions fitted to the FRAP data acquired from *n* cells (shown in the plots). For single-cell FRAP data and additional quantification, see Supplementary Fig. 6d-k. **d**, DPPA3-mediated inhibition of UHRF1 chromatin binding is necessary and sufficient to promote DNA demethylation. Percentage of DNA methylation change at LINE-1 elements (%) in D3KO ESCs after induction of the indicated mutant forms of *Dppa3* as measured by bisulfite sequencing of *n* = 4 biological replicates. In the boxplot in **c**, horizontal black lines within boxes represent median values, boxes indicate the upper and lower quartiles, whiskers indicate the 1.5 interquartile range, and dots indicate outliers. P-values based on Welch’s two-sided t-test

Nevertheless, DPPA3-ΔNES and DPPA3 61-150 managed to significantly disrupt the focal patterning and association with chromocenters of UHRF1 within the nucleus itself, with DPPA3-ΔNES causing an even greater reduction in the CV of nuclear UHRF1 than DPPA3-WT (Fig. 5b and Supplementary Fig. 6c). In contrast, the loss or mutation of residues critical for its interaction with UHRF1 compromised DPPA3’s ability to effectively restore the diffuse localization of nuclear UHRF1 (Fig. 5b and Supplementary Fig. 6c). On the one hand, FRAP analysis revealed that the disruption or deletion of the UHRF1 interaction interface (DPPA3-KRR, DPPA3-R107E, DPPA3 1-60) severely diminished the ability of DPPA3 to release UHRF1 from chromatin (Fig. 5c and Supplementary Fig. 6f-k). On the other hand, the C-terminal half of DPPA3 lacking nuclear export signal but retaining UHRF1 interaction came close to fully restoring the mobility of UHRF1 (Fig. 5c and Supplementary Fig. 6i-k). Remarkably, DPPA3-ΔNES mobilized UHRF1 to an even greater extent than DPPA3-WT (Fig. 5c and Supplementary Fig. 6d,e,j,k), suggesting that operative nuclear export might even antagonize DPPA3-mediated inhibition of UHRF1 chromatin binding. Supporting this notion, chemical inhibition of nuclear export using leptomycin-B (LMB) significantly enhanced the inhibition of UHRF1 chromatin binding in U1G/D3KO ESCs expressing DPPA3-WT (Supplementary Fig. 5h-k). Taken together our data shows that the efficiency of DPPA3-dependent release of UHRF1 from chromatin requires its interaction with UHRF1 but not its nuclear export.

To further address the question whether the nucleocytoplasmic translocation of UHRF1 and impaired UHRF1 chromatin binding both contribute to DPPA3-mediated inhibition of DNA methylation maintenance, we assessed the ability of each DPPA3 mutant to rescue the hypermethylation of LINE-1 elements in Dppa3KO ESCs (Fig. 5d). Strikingly, DPPA3-ΔNES fully rescued the hypermethylation and achieved a greater loss of DNA methylation than DPPA3-WT, whereas DPPA3 mutants lacking the residues important for UHRF1 binding failed to restore low methylation levels (Fig. 5d). Overall, the ability of each DPPA3 mutant to reduce DNA methylation levels closely mirrored the extent to which each mutant impaired UHRF1 chromatin binding (Fig. 5c and Supplementary Fig. 6d-k). In line with the increased mobility of UHRF1 occuring in the absence of DPPA3 nuclear export (Fig. 5c, Supplementary Figs. 5h-k and 6d,e,j,k), the nucleocytoplasmic translocation of UHRF1 is not only dispensable but rather attenuates DPPA3-mediated inhibition of maintenance methylation (Fig. 5d). Collectively, our findings demonstrate the inhibition of UHRF1 chromatin binding, as opposed to nucleocytosolic translocation of UHRF1, to be the primary mechanism by which DPPA3 drives hypomethylation in naïve ESCs.

### DPPA3 binds nuclear UHRF1 with high affinity prompting its release from chromatin in ESCs

We next set out to investigate the mechanistic basis of DPPA3’s ability to inhibit UHRF1 chromatin binding in naïve ESCs. DPPA3 has been reported to specifically bind H3K9me2^61^, a histone modification critical for UHRF1 targeting^68,75,76^. These prior findings led us to consider two possible mechanistic explanations for DPPA3-mediated UHRF1 inhibition in naïve ESCs: 1) DPPA3 blocks access of UHRF1 to chromatin by competing in binding to H3K9me2, 2) DPPA3 directly or indirectly binds to UHRF1 and thereby prevents it from accessing chromatin.

To simultaneously assess the dynamics of both UHRF1 and DPPA3 under physiological conditions in live ES cells, we employed raster image correlation spectroscopy with pulsed interleaved excitation (PIE-RICS) (Fig. 6a). RICS is a confocal imaging method which enables the measurement of binding and diffusive properties in living cells. Using images acquired on a laser scanning confocal microscope, spatiotemporal information of fluorescently labeled proteins can be extracted from the shape of the spatial autocorrelation function (ACF). A diffusive model is fitted to the ACF which yields the average diffusion coefficient, the concentration, and the fraction of bound molecules^77^. If two proteins are labeled with distinct fluorophores and imaged simultaneously with separate detectors, the extent of their interaction can be extracted from the cross-correlation of their fluctuations using cross-correlation RICS (ccRICS) (Fig. 6a)^78^.

**Fig. 6:**
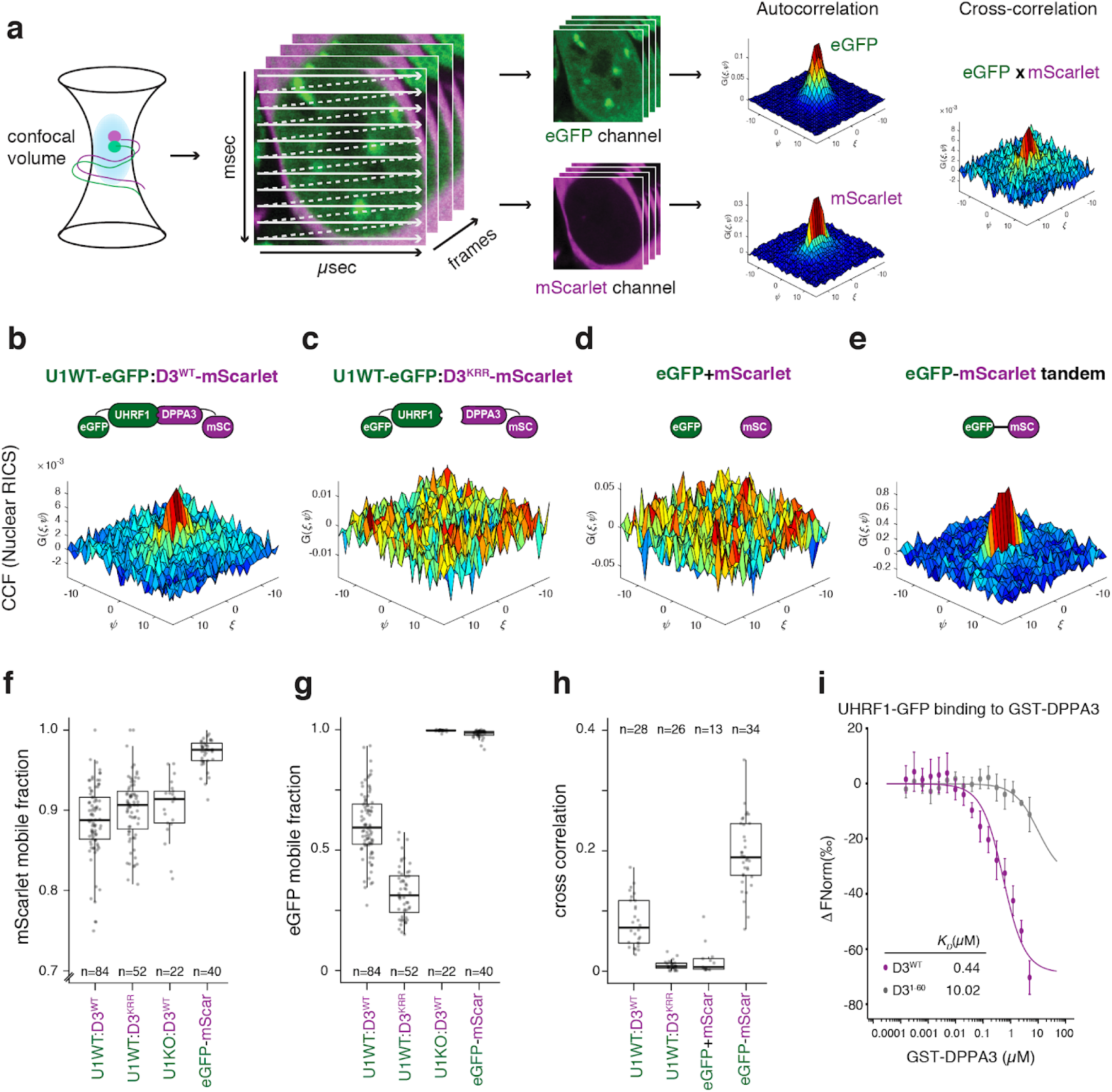
DPPA3 binds nuclear UHRF1 with high affinity prompting its release from chromatin in ESCs. **a**, Overview of RICS and ccRICS. Confocal image series are acquired using a calibrated point-scanning laser, generating spatio-temporal fluorescence information on the microsecond and millisecond timescales. An autocorrelation function (ACF) is calculated from the fluorescence fluctuations and used to fit a diffusive model^77^. The cross-correlation of fluctuations between two channels is used to estimate the co-occurence of two fluorescent molecules within live cells^78^. The mean cross-correlation of fluctuations is calculated and shown in the 3D plot, color-coded according to the correlation value. **b,e**, Representative plots of the cross-correlation function (CCF) between the depicted fluorescent molecules in cells from each cell line measured, including U1G/D3KO + pSBtet-D3 ESCs expressing the following forms of DPPA3-mScalet: (**b**) wild-type (U1WT:D3^WT^) and (**c**) K85E/R85E/K87E mutant (U1WT:D3^KRR^), and control ESCs expressing (**d**) free eGFP and free mScarlet (eGFP + mScarlet) and (**e**) an eGFP-mScarlet tandem fusion (eGFP-mScarlet). See Supplementary Fig. 7 for the images and ACF plots of the cells used to make the representative CCF plots. **f,g**, Mobile fraction of (**f**) mScarlet and (**g**) eGFP species in the cell lines depicted in (b, c, and e) as well as in Uhrf1KO ESCs expressing free eGFP and wild-type DPPA3-mScarlet (U1KO:D3^WT^) (Supplementary Fig. 7c). The mobile fraction was derived from a two-component model fit of the autocorrelation function. Data are pooled from three (U1WT:D3^WT^, U1WT:D3^KRR^) or two (U1KO:D3^WT^, eGFP-mScar) independent experiments. **h**, Mean cross-correlation values of mobile eGFP and mScarlet measured in the cell lines depicted in (**b-e**). The fast timescale axis is indicated by *ξ*, and the slow timescale axis is indicated by *ψ*. Data are pooled from two independent experiments.**i**, Microscale thermophoresis measurements of UHRF1-eGFP binding to GST-DPPA3 WT (D3^WT^) or GST-DPPA3 1-60 (D3^1-60^). Error bars indicate mean ± SEM of *n* = 2 technical replicates from *n* = 4 independent experiments. In **f-h**, each data point represents the measured and fit values from a single cell where *n* = number of cells measured (indicated in the plots). In the box plots, darker horizontal lines within boxes represent median values. The limits of the boxes indicate upper and lower quartiles, and whiskers indicate the 1.5-fold interquartile range.

We first measured the mobility of DPPA3-mScarlet variants expressed in U1GFP/D3KOs (Supplementary Fig. 7a,b). RICS analysis revealed that over the timescale of the measurements, nuclear DPPA3-WT was predominantly unbound from chromatin and freely diffusing through the nucleus at a rate of 7.18 ± 1.87 µm^2^/s (Supplementary Fig. 7f). The fraction of mobile DPPA3-mScarlet molecules was measured to be 88.4 ± 5.2% (Fig. 6f), validating globally weak binding inferred from ChIP-Seq profiles^60^. These mobility parameters were largely unaffected by disruption of the UHRF1 interaction, with the DPPA3-KRR mutant behaving similarly to wild-type DPPA3 (Fig. 6f and Supplementary Fig. 7f). To rule out a potential competition between UHRF1 and DPPA3 for H3K9me2 binding, we next used RICS to determine if DPPA3 dynamics are altered in the absence of UHRF1. For this purpose, we introduced the DPPA3-WT-mScarlet cassette into Uhrf1KO (U1KO) ESCs^79^, in which free eGFP is expressed from the endogenous *Uhrf1* promoter (Supplementary Fig. 7c). However, neither the diffusion rate nor the mobile fraction of DPPA3 were appreciably altered in cells devoid of UHRF1, suggesting the high fraction of unbound DPPA3 to be unrelated to the presence of UHRF1 (Fig. 6f and Supplementary Fig. 7f). Overall, our RICS data demonstrate that, in contrast to zygotes ^61^, DPPA3 in ESCs lacks a strong capacity for chromatin binding, and as such is not engaged in competition with UHRF1 for chromatin binding.

We next used RICS to analyse the dynamics of UHRF1-GFP in response to DPPA3 induction (Fig. 6a). In cells expressing DPPA3-KRR, RICS measurements revealed that only 32.4 ± 10% of UHRF1 is mobile, indicating that the majority of UHRF1 is chromatin-bound (Fig. 6g). In contrast, expression of wild-type DPPA3 lead to a dramatic increase in the mobile fraction UHRF1 (60.6 ± 13.7% mobile fraction for UHRF1) (Fig. 6g and Supplementary Fig. 7g,h). Furthermore, the mobile fraction of UHRF1 increased as a function of the relative abundance of nuclear DPPA3 to UHRF1 (Supplementary Fig. 7i), thereby indicating a stoichiometric effect of DPPA3 on UHRF1 chromatin binding, consistent with a physical interaction. Thus, these results demonstrate that DPPA3 potently disrupts UHRF1 chromatin binding in live ESCs and suggest its interaction with UHRF1 to be critical to do so.

To determine whether such an interaction is indeed present in the nuclei of live ESCs, we performed cross-correlation RICS (ccRICS) (Fig. 6a). We first validated ccRICS in ESCs by analyzing live cells expressing a tandem eGFP-mScarlet fusion (Fig. 6e and Supplementary Fig. 7d), or expressing both freely diffusing eGFP and mScarlet (Fig. 6d and Supplementary Fig. 7e). For the tandem eGFP-mScarlet fusion, we observed a clear positive cross-correlation indicative of eGFP and mScarlet existing in the same complex (Fig. 6e,h), as would be expected for an eGFP-mScarlet fusion. On the other hand, freely diffusing eGFP and mScarlet yielded no visible cross-correlation (Fig. 6d,h), consistent with two independent proteins which do not interact. Upon applying ccRICS to nuclear UHRF1-GFP and DPPA3-mScarlet, we observed a prominent cross-correlation between wild-type DPPA3 and the primarily unbound fraction of UHRF1 (Fig. 6b,h), indicating that mobilized UHRF1 exists in a high affinity complex with DPPA3 in live ESCs. In marked contrast, DPPA3-KRR and UHRF1-GFP failed to exhibit detectable cross-correlation (Fig. 6c,h), consistent with the DPPA3-KRR mutant’s diminished capacity to bind^49^ and mobilize UHRF1 (Fig. 5c and Supplementary Fig. 6f,j,k). Overall, these findings demonstrate that nuclear DPPA3 interacts with UHRF1 to form a highly mobile complex in naïve ESCs which precludes UHRF1 chromatin binding.

To determine whether the DPPA3-UHRF1 complex identified *in vivo* (Fig. 6h) corresponds to a high affinity direct interaction, we performed microscale thermophoresis (MST) measurements using recombinant UHRF1-GFP and DPPA3 proteins. MST analysis revealed a direct and high affinity (K_D_: 0.44 µM) interaction between the DPPA3 WT and UHRF1 (Fig. 6i). No binding was observed for DPPA3 1-60, lacking the residues essential for interaction with UHRF1 (Fig. 6i). In line with the results obtained by ccRICS, these data support the notion that DPPA3 directly binds UHRF1 *in vivo*. Interestingly, the affinity of the UHRF1-DPPA3 interaction was comparable or even far greater than that reported for the binding of UHRF1 to H3K9me3 or unmodified H3 peptides, respectively^80,81^.

To better understand how UHRF1 chromatin loading is impaired by its direct interaction with DPPA3, we applied a fluorescent-three-hybrid (F3H) assay to identify the UHRF1 domain bound by DPPA3 *in vivo* (Supplementary Fig. 7j,k). In short, this method relies on a cell line harboring an array of lac operator binding sites in the nucleus at which a GFP-tagged “bait” protein can be immobilized and visualized as a spot. Thus, the extent of recruitment of an mScarlet-tagged “prey” protein to the nuclear GFP-spot offers a quantifiable measure of the interaction propensity of the “bait” and “prey” proteins *in vivo* (Supplementary Fig. 7k)^82^. Using UHRF1-GFP domain deletions as the immobilized bait (Supplementary Fig. 7j,k), we assessed how the loss of each domain affected the recruitment of mDPPA3-mScarlet to the GFP spot. In contrast to the other UHRF1 domain deletions, removal of the PHD domain essentially abolished recruitment of DPPA3 to the lac spot, demonstrating DPPA3 binds UHRF1 via its PHD domain *in vivo* (Supplementary Fig. 7l,m). The PHD of UHRF1 is essential for its recruitment to chromatin^73,81,83^, ubiquitination of H3 and recruitment of DNMT1 to replication foci^66,67^. Thus, our *in vivo* results suggest that the high affinity interaction of DPPA3 with UHRF1’s PHD domain precludes UHRF1 from binding chromatin in ESCs, which is also supported by a recent report demonstrating that DPPA3 specifically binds the PHD domain of UHRF1 to competitively inhibit H3 tail binding *in vitro*^84^.

### DPPA3 can inhibit UHRF1 function and drive global DNA demethylation in distantly related, non-mammalian species

Whereas UHRF1 and TET proteins are widely conserved throughout plants and vertebrates^34,85^, both early embryonic global hypomethylation^86^ and the *Dppa3* gene are unique to mammals. Consistent with UHRF1’s conserved role in maintenance DNA methylation, a multiple sequence alignment of UHRF1’s PHD domain showed that the residues critical for the recognition of histone H3 are completely conserved from mammals to invertebrates (Fig. 7a). This prompted us to consider the possibility that DPPA3 might be capable of modulating the function of distantly related UHRF1 homologs outside of mammals. To test this hypothesis, we used amphibian (*Xenopus laevis*) egg extracts to assess the ability of mouse DPPA3 (mDPPA3) to interact with a non-mammalian form of UHRF1. Despite the 360 million year evolutionary distance between mouse and *Xenopus*^87^, mDPPA3 not only bound *Xenopus* UHRF1 (xUHRF1) with high affinity (Fig. 7b,c and Supplementary Fig. 8a,b) it also interacted with xUHRF1 specifically via its PHD domain (Supplementary Fig. 8c-e). Moreover, the first 60 amino acids of DPPA3 were dispensable for its interaction with UHRF1 (Supplementary Fig. 8a,b). Interestingly, mutation to R107, reported to be critical for DPPA3’s binding with mouse UHRF1^49^, diminished but did not fully disrupt the interaction (Supplementary Fig. 8b,e). The R107E mutant retained the ability to bind the xUHRF1-PHD domain but exhibited decreased binding to xUHRF1-PHD-SRA under high-salt conditions (Supplementary Fig. 8e), suggesting that R107E changes the binding mode of mDPPA3 to xUHRF1, rather than inhibiting the complex formation. Considering the remarkable similarity between DPPA3’s interaction with mouse and *Xenopus* UHRF1, we reasoned that the ability of DPPA3 to inhibit UHRF1 chromatin binding and maintenance DNA methylation might be transferable to *Xenopus*. To address this, we took advantage of a cell-free system derived from interphase *Xenopus* egg extracts to reconstitute DNA maintenance methylation^66^. Remarkably, recombinant mDPPA3 completely disrupted chromatin binding of both *Xenopus* UHRF1 and DNMT1 without affecting the loading of replication factors such as xCDC45, xRPA2, and xPCNA (Fig. 7d). We determined that the inhibition of xUHRF1 and xDNMT1 chromatin loading only requires DPPA3’s C-terminus (61-150 a.a.) and is no longer possible upon mutation of R107 (R107E) (Supplementary Fig. 8h), in line with our results in mouse ESCs (Fig. 5d). Moreover, DPPA3-mediated inhibition of xUHRF1 chromatin loading resulted in the severe perturbation of histone H3 dual-monoubiquitylation (H3Ub2), which is necessary for the recruitment of DNMT1^66,67,88^ (Supplementary Fig. 8f). To determine whether mDPPA3 can displace xUHRF1 already bound to chromatin, we first depleted *Xenopus* egg extracts of xDNMT1 to stimulate the hyper-accumulation of xUHRF1 on chromatin^66,89^ and then added recombinant mDPPA3 after S-phase had commenced (Supplementary Fig. 8g). Under these conditions, both wild-type mDPPA3 and the 61-150 fragment potently displaced xUHRF1 from chromatin, leading to suppressed H3 ubiquitylation (Supplementary Fig. 8g).

**Fig. 7:**
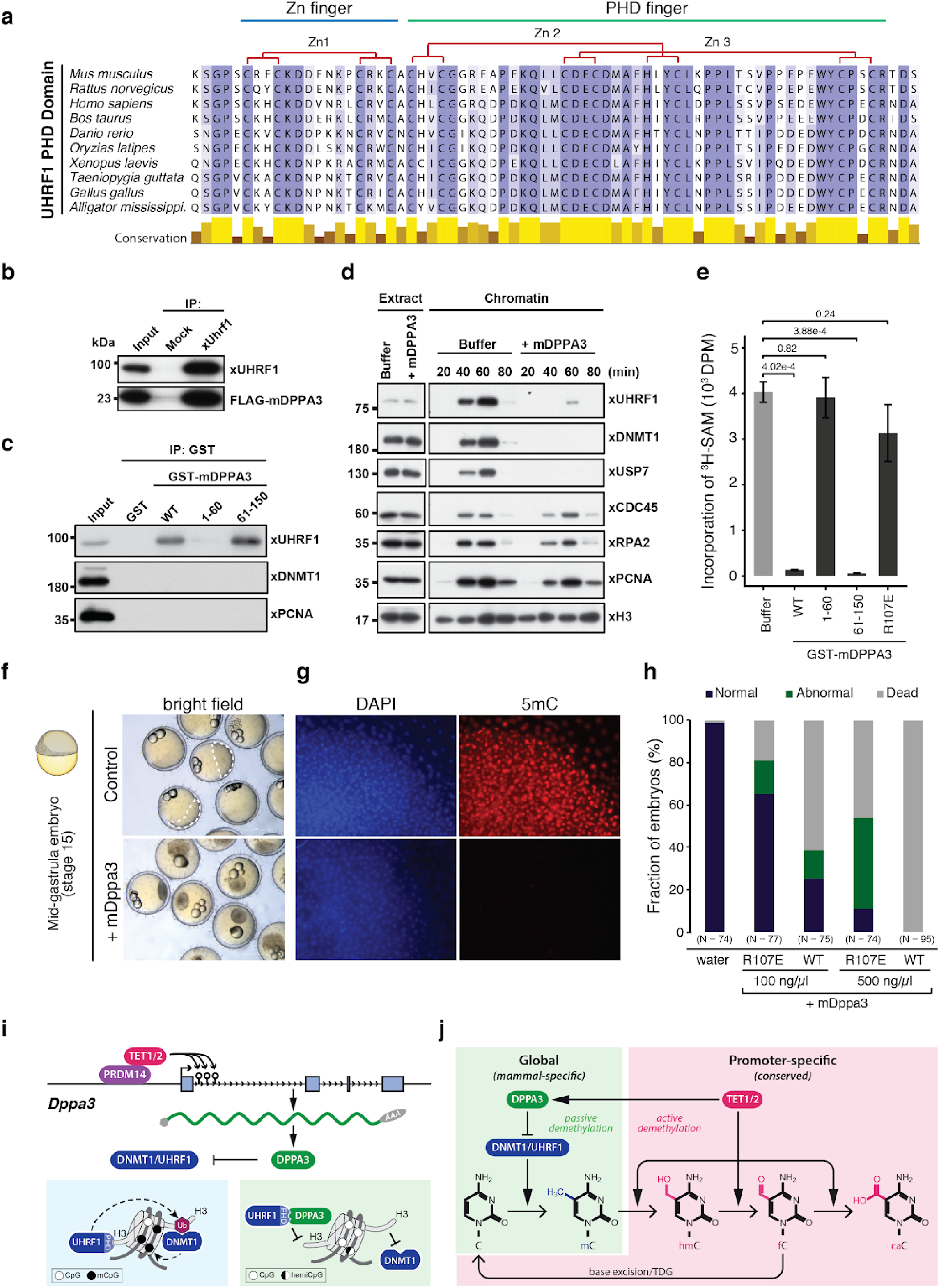
DPPA3 evolved in boreoeutherian mammals but also functions in lower vertebrates. **a**, Protein sequence alignment of the PHD domain of the UHRF1 family. The UHRF1 PHD domain shows high amino acid conservation throughout vertebrates, especially the residues involved in Zinc coordination (indicated above). **b**, Endogenous xUHRF1 binds mDPPA3. IPs were performed on *Xenopus* egg extracts incubated with FLAG-mDPPA3 using either a control (Mock) or anti-xUHRF antibody and then analyzed by immunoblotting using the indicated antibodies. **c**, xUHRF1 binds the C-terminus of mDPPA3. GST-tagged mDPPA3 wild-type (WT), point mutant R107E, and truncations (1-60 and 61-150) were immobilized on GSH beads and incubated with *Xenopus* egg extracts. Bound proteins were analyzed using the indicated antibodies. **d**, mDPPA3 inhibits xUHRF1 and xDNMT1 chromatin binding. Sperm chromatin was incubated with interphase *Xenopus* egg extracts supplemented with buffer (+buffer) or GST-mDPPA3 (+mDPPA3). Chromatin fractions were isolated and subjected to immunoblotting using the antibodies indicated. **e**, mDPPA3 inhibits maintenance DNA methylation in *Xenopus*. The efficiency of maintenance DNA methylation was assessed by the incorporation of radiolabelled methyl groups from S-[methyl-^3^H]-adenosyl-L-methionine (^3^H-SAM) into DNA purified from egg extracts. Disintegrations per minute (DPM). Error bars indicate mean ± SD calculated from *n* = 4 independent experiments. Depicted p-values based on a Student’s two-sided t-test. **f**, mDPPA3 compromises the embryonic development of medaka. Representative images of developing mid-gastrula stage embryos (control injection) and arrested, blastula stage embryos injected with *mDppa3*. Injections were performed on one-cell stage embryos and images were acquired ∼18 h after fertilization. **g**, mDPPA3 drives global DNA demethylation in medaka embryos. Representative 5mC immunostainings in control and *mDppa3*-injected medaka embryos at the late blastula stage (∼8 h after fertilization). The assay was repeated > 3 times with similar results. Scale bars represent 50 µm. DNA counterstain: DAPI,4′,6-diamidino-2-phenylindole. **h**, Global hypomethylation causes developmental arrest in medaka. Percentage of normal, abnormal, or dead. Embryos were injected with wild-type *mDppa3* (WT) or *mDppa3* R107E (R107E) at two different concentrations (100 ng/µl or 500 ng/µl) or water at the one-cell stage and analyzed ∼18 h after fertilization. N = number of embryos from *n* = 3 independent injection experiments. **i**, TET1 and TET2 are recruited by PRDM14 to the promoter of *Dppa3* where they promoter active DNA demethylation and transcription of *Dppa3*. DPPA3 is expressed and inhibits maintenance DNA methylation by directly binding UHRF1 and releasing it from chromatin. **j**, TET1 and TET2 control DNA methylation levels by two evolutionary and mechanistically distinct pathways.

We next assessed the effect of DPPA3 on *Xenopus* maintenance DNA methylation. Consistent with the severe disruption of xDNMT1 chromatin loading, both DPPA3 wild-type and 61-150 effectively abolished replication-dependent DNA methylation in *Xenopus* egg extracts (Fig. 7e). In contrast, DPPA3 1-60 and DPPA3 R107E, which both failed to suppress xUHRF1 and xDNMT1 binding, did not significantly alter maintenance DNA methylation activity (Fig. 7e and Supplementary Fig. 8d,e). Taken together, our data demonstrate DPPA3 to be capable to potently inhibit maintenance DNA methylation in a non-mammalian system.

These findings raised the question whether a single protein capable of inhibiting UHRF1 function like DPPA3 could establish a mammalian-like global hypomethylation during the early embryonic development of a non-mammalian organism. To explore this possibility we turned to the biomedical model fish, medaka (*Oryzias latipes*), which does not exhibit genome-wide erasure of DNA methylation^90^ and diverged from mammals 450 million years ago^87^. We injected medaka embryos with *Dppa3* mRNA at the one-cell stage and then tracked their developmental progression. Remarkably, medaka embryos injected with *Dppa3* failed to develop beyond the blastula stage (Fig. 7f) and exhibited a near-complete elimination of global DNA methylation as assessed by immunofluorescence (Fig. 7g). DPPA3-mediated DNA methylation loss was both dose dependent and sensitive to the R107E mutation, which induced only partial demethylation (Supplementary Fig. 8i). Interestingly, medaka embryos injected with DPPA3 R107E showed far fewer developmental defects than those injected with wild-type DPPA3 (Fig. 7h), suggesting that the embryonic arrest resulting from DPPA3 expression is a consequence of the global loss of DNA methylation. Taken together, these results demonstrate that mammalian DPPA3 can inhibit UHRF1 to drive passive demethylation in distant, non-mammalian contexts.

## DISCUSSION

While the appearance of genome-wide DNA demethylation in mammals represents a momentous change with far-reaching consequences to epigenetic gene regulation during early development, the key enzymes involved in DNA modification are highly conserved in vertebrates. As the role of TET enzymes in active demethylation is well documented^91^, we investigated their contribution to the hypomethylated state of naïve ESCs. Mutation of the catalytic core of TET enzymes caused - as expected - a genome-wide increase in DNA methylation but mostly at sites where TET proteins do not bind suggesting a rather indirect mechanism. Among the few genes depending on TET activity for expression in naïve ESCs and downregulated at the transition to EpiLCs was *Dppa3*. Demethylation at the *Dppa3* locus coincides with TET1 and TET2 binding and TDG dependent removal of oxidized cytosine residues via base excision repair. DPPA3 in turn binds and displaces UHRF1 from chromatin and thereby prevents the recruitment of DNMT1 and the maintenance of DNA methylation in ESCs (see graphic summary in Fig. 7).

Despite long recognized as a marker of naïve ESCs resembling the inner cell mass^53,92^, we provide, to our knowledge, the first evidence that DPPA3 directly promotes the genome-wide DNA hypomethylation characteristic of mammalian naïve pluripotency. This unique pathway, in which TET proteins indirectly cause passive demethylation, is based upon two uniquely mammalian innovations: the expression of TET genes in pluripotent cell types^39,63,93^ and the evolution of the novel *Dppa3* gene, positioned within a pluripotency gene cluster and dependent on TET activity for expression. In support of this novel pathway for passive demethylation, we found that TET mutant ESCs show a similar phenotype as Dppa3KO cells with respect to UHRF1 inhibition and hypomethylation and can be rescued by ectopic expression of *Dppa3*.

Our findings also provide the missing link to reconcile previous, apparently conflicting reports. To date, three distinct mechanisms have been proposed for global hypomethylation accompanying naïve pluripotency: TET-mediated active demethylation^37,40,94^, impaired maintenance DNA methylation^94^, and PRDM14-dependent suppression of methylation^36,37,51^. As both a downstream target of TETs and PRDM14 as well as a direct inhibitor of maintenance DNA methylation, DPPA3 mechanistically links and integrates these three proposed pathways of demethylation (see graphic summary in Fig. 7).

Our mechanistic data showing DPPA3 to displace UHRF1 and DNMT1 from chromatin provide a conclusive explanation for the previous observation that global hypomethylation in naïve ESCs was accompanied by reduced levels of UHRF1 at replication foci^94^. The hypomethylated state of naïve ESCs has also been reported to be dependent on PRDM14^36,51^, which has been suggested to promote demethylation by repressing *de novo* DNA methyltransferases^36,40,51,57^. However, recent studies have demonstrated that the loss of *de novo* methylation only marginally affects DNA methylation levels in mouse and human ESCs^94,95^. Interestingly, the loss of *Prdm14* lead to global hypermethylation and, to our surprise, also the downregulation of *Dppa3*^51,96,97^. Our results suggest that the reported ability of PRDM14 to promote hypomethylation in naïve ESCs largely relies on its activation of the *Dppa3* gene ultimately leading to an inhibition of maintenance methylation.

The comparison of TET catalytic mutants and Dppa3KO ESCs allows us to distinguish TET-dependent passive DNA demethylation mediated by DPPA3 from *bona fide* active demethylation. We show that TET activity is indispensable for the active demethylation of a subset of promoters in naïve ESCs, especially those of developmental genes. These findings uncover two evolutionary and mechanistically distinct functions of TET catalytic activity. Whereas TET-mediated active demethylation of developmental genes is evolutionarily conserved among vertebrates^63,98–100^, the use of TET proteins to promote global demethylation appears to be specific to mammalian pluripotency^37,40,94^ and mediated by the recently evolved *Dppa3* (Figs. 2c and 7j).

To date, our understanding of DPPA3’s function in the regulation of DNA methylation had been clouded by seemingly conflicting reports from different developmental stages and cell types. DPPA3’s ability to modulate DNA methylation was first described in the context of zygotes^48^, where it was subsequently demonstrated to specifically protect the maternal genome from TET3-dependent demethylation^27,58,101^. In contrast, DPPA3 was later shown to facilitate DNA demethylation during PGC specification^46^, iPSC reprogramming^59^ and oocyte maturation^102,103^. Whereas DPPA3 was shown to disrupt UHRF1 function by sequestering it to the cytoplasm in oocytes^102^, we demonstrate that DPPA3-mediated nucleocytoplasmic translocation of UHRF1 is not only dispensable but actually attenuates DPPA3’s promotion of hypomethylation in ESCs. In light of our data from naïve ESCs, *Xenopus*, and medaka, DPPA3’s capacity to directly bind UHRF1’s PHD domain and thereby inhibit UHRF1 chromatin binding appears to be its most basal function. Considering that DPPA3 localization is highly dynamic during the different developmental time periods at which it is expressed^46,62,104^, it stands to reason that its role in modulating DNA methylation might also be dynamically regulated by yet-to-be determined regulatory mechanisms. For example, immediately following fertilization, full length DPPA3 is cleaved and its C-terminal domain is specifically degraded^62^. Interestingly, we identified this exact C-terminal stretch of DPPA3 to be necessary and sufficient for DPPA3’s inhibition of maintenance DNA methylation. Thus, the precisely timed destruction of this crucial domain might offer an explanation for the differing roles of DPPA3 in regulating DNA methylation between oocytes and zygotes^58,101–103^.

As the most basic and evolutionarily conserved function of DNA methylation is the repression of transposable elements^105^, the emergence of genome-wide DNA demethylation in mammals raises several fundamental questions. While the DPPA3 mediated erasure of parental DNA methylation might facilitate the establishment of new epigenetic patterns during development, it should be noted that non-mammalian vertebrates manage to undergo normal development without genome-wide demethylation. Moreover, the global loss of DNA methylation even poses a severe threat as excessive demethylation triggers derepression of TEs leading to genomic instability and ultimately cell death in most cell types^7–12^. Remarkably, mammalian naïve pluripotent cell types seem to have acquired the ability to tolerate global hypomethylation, suggesting that the evolutionary emergence of DPPA3 was likely accompanied by measures to control and productively integrate this new factor in epigenetic regulation in ESCs. In fact, many TEs in mammals are not only expressed during development but appear to have been co-opted to drive transcriptional networks critical for the establishment of pluripotency and progression through pre-implantation development^106,107^.

A good example for the functional integration of TEs in regulatory networks is the reactivation of endogenous retroviruses (ERVs) that is critical for the maternal-to-zygotic transition (MZT), the first major developmental step when maternal mRNAs are degraded and zygotic transcription begins^108–111^. Strikingly, the activation of ERVs is severely impaired in *Dppa3* knockout embryos resulting in MZT failure^60^. It is tempting to speculate that mammal-specific demethylation originates from an arms race between TE and host. DPPA3 may have arisen as a means to overcome the host defence system and was then co-opted by the host and gradually integrated into regulatory networks during evolution. Such a scenario is compatible with the unique occurrence of *Dppa3* in mammals and with our finding that DPPA3 alone is sufficient to inhibit DNA methylation maintenance in *Xenopus* and medaka, species that harbor no *Dppa3* gene and exhibits constant DNA methylation levels at all stages of development^13,14^. Follow-up studies that investigate the origin of *Dppa3* and whether a similar rewiring of early development may have occurred in other, not yet studied branches of vertebrates, are needed to further explore this evolutionary scenario.

## MATERIALS AND METHODS

### Cell culture

Naïve J1 mouse ESCs were cultured and differentiated into EpiLCs as described previously^112,113^. In brief, for both naïve ESCs and EpiLCs defined media was used, consisting of N2B27: 50% neurobasal medium (Life Technologies), 50% DMEM/F12 (Life Technologies), 2 mM L-glutamine (Life Technologies), 0.1 mM β-mercaptoethanol (Life Technologies), N2 supplement (Life Technologies), B27 serum-free supplement (Life Technologies), 100 U/mL penicillin, and 100 μg/mL streptomycin (Sigma). Naïve ESCs were maintained on flasks treated with 0.2% gelatin in defined media containing 2i (1 μM PD032591 and 3 μM CHIR99021 (Axon Medchem, Netherlands)), 1000 U/mL recombinant leukemia inhibitory factor (LIF, Millipore), and 0.3% BSA (Gibco) for at least three passages before commencing differentiation. For reprogramming experiments, naïve media was supplemented with freshly prepared 100 µM Vitamin C (L-ascorbic acid 2-phosphate, Sigma). To differentiate naïve ESCs into Epiblast-like cells (EpiLCs), flasks were first pre-treated with Geltrex (Life Technologies) diluted 1:100 in DMEM/F12 (Life Technologies) and incubated at 37 °C overnight. Naïve ESCs were plated on Geltrex-treated flasks in defined medium containing 10 ng/mL Fgf2 (R&D Systems), 20 ng/mL Activin A (R&D Systems) and 0.1× Knockout Serum Replacement (KSR) (Life Technologies). Media was changed after 24 h and EpiLCs were harvested for RRBS and RNA-seq experiments after 48 h.

For CRISPR-assisted cell line generation, mouse ESCs were maintained on 0.2% gelatin-coated dishes in Dulbecco’s modified Eagle’s medium (Sigma) supplemented with 16% fetal bovine serum (FBS, Sigma), 0.1 mM ß-mercaptoethanol (Invitrogen), 2 mM L-glutamine (Sigma), 1× MEM Non-essential amino acids (Sigma), 100 U/mL penicillin, 100 μg/mL streptomycin (Sigma), homemade recombinant LIF tested for efficient self-renewal maintenance, and 2i (1 μM PD032591 and 3 μM CHIR99021 (Axon Medchem, Netherlands)).

For experiments in which cells were propagated in “serum LIF” conditions, the cells were maintained on 0.2% gelatin-coated dishes in Dulbecco’s modified Eagle’s medium (Sigma) supplemented with 16% fetal bovine serum (FBS, Sigma), 0.1 mM ß-mercaptoethanol (Invitrogen), 2 mM L-glutamine (Sigma), 1× MEM Non-essential amino acids (Sigma), 100 U/mL penicillin, 100 μg/mL streptomycin (Sigma), LIF (ESGRO, Millipore).

HESCs (line H9) were maintained in mTeSR1 medium (05850, STEMCELL Technologies) on Matrigel-coated plates (356234, Corning) prepared by 1:100 dilution, and 5 ml coating of 10 cm plates for 1 h at 37 °C. Colonies were passaged using the gentle cell dissociation reagent (07174, StemCell Technologies). All cell lines were regularly tested for Mycoplasma contamination by PCR.

### Sleeping Beauty Constructs

To generate the sleeping beauty donor vector with an N-terminal 3xFLAG tag and a fluorescent readout of doxycycline induction, we first used primers with overhangs harboring SfiI sites to amplify the IRES-DsRed-Express from pIRES2-DsRed-Express (Clontech). This fragment was then cloned into the NruI site in pUC57-GentR via cut-ligation to generate an intermediate cloning vector pUC57-SfiI-IRES-DsRed-Express-SfiI. A synthesized gBlock (IDT, Coralville, IA, USA) containing Kozak-BIO-3XFLAG-AsiSI-NotI-V5 was cloned into the Eco47III site of the intermediate cloning vector via cut-ligation. The luciferase insert from pSBtet-Pur^114^ (Addgene plasmid #60507) was excised using SfiI. The SfiI-flanked Kozak-BIO-3XFLAG-AsiSI-NotI-V5-IRES-DsRed-Express cassette was digested out of the intermediate cloning vector using SfiI and ligated into the pSBtet-Pur vector backbone linearized by SfiI. The end result was the parental vector, pSBtet-3xFLAG-IRES-DsRed-Express-PuroR. The pSBtet-3x-FLAG-mScarlet-PuroR vector was constructed by inserting a synthesized gBlock (IDT, Coralville, IA, USA) containing the SfiI-BIO-3XFLAG-AsiSI-NotI-mScarlet sequence into the SfiI-linearized pSBtet-Pur vector backbone using Gibson assembly^115^. For *Dppa3* expression constructs, the coding sequence of wild-type and mutant forms of *Dppa3* were synthesized as gBlocks (IDT, Coralville, IA, USA) and inserted into the pSBtet-3xFLAG-IRES-DsRed-Express-PuroR vector (linearized by AsiSI and NotI) using Gibson assembly. To produce the *Dppa3*-mScarlet fusion expression constructs, wild-type and mutant forms of *Dppa3* were amplified from pSBtet-3xFLAG-Dppa3-IRES-DsRed-Express-PuroR constructs using primers with overhangs homologous to the AsiSI and NotI restriction sites of the pSBtet-3x-FLAG-mScarlet-PuroR vector. Wild-type and mutant *Dppa3* amplicons were subcloned into the pSBtet-3x-FLAG-mScarlet-PuroR vector (linearized with AsiSI and NotI) using Gibson assembly.

For experiments involving the SBtet-3xFLAG-*Dppa3* cassette, all inductions were performed using 1 µg/mL doxycycline (Sigma-Aldrich). The DPPA3-WT construct was able to rescue the cytoplasmic localization and chromatin association of UHRF1 indicating that C-terminally tagged DPPA3 remains functional (Fig. 5b-d).

### CRISPR/Cas9 genome engineering

For the generation of *Tet1, Tet2*, and *Tet1*/*Tet2* catalytic mutants, specific gRNAs targeting the catalytic center of *Tet1* and *Tet2* were cloned into a modified version of the SpCas9-T2A-GFP/gRNA (px458^116^, Addgene plasmid #48138), to which we fused a truncated form of human Geminin (hGem) to SpCas9 in order to increase homology-directed repair efficiency^117^.

A 200 bp ssDNA oligonucleotide harboring the H1652Y and D1654A mutations and ∼100 bp of homology to the genomic locus was synthesized (IDT, Coralville, IA, USA). For targetings in wild-type J1 ESCs, cells were transfected with a 4:1 ratio of donor oligo and Cas9/gRNA construct. Positively transfected cells were isolated based on GFP expression using fluorescence-activated cell sorting (FACS) and plated at clonal density in ESC media 2 days after transfection. Cell lysis in 96-well plates, PCR on lysates, and restriction digests were performed as previously described^113^. *Tet1* catalytic mutation was confirmed by Sanger sequencing.

As C-terminally tagged GFP labeled UHRF1 transgenes were shown to be able to rescue U1KO^67^, the tagging of endogenous *Uhrf1* was also performed at the C-terminus. For insertion of the HALO or eGFP coding sequence into the endogenous *Dppa3* and *Uhrf1* loci, respectively, *Dppa3* and *Uhrf1* specific gRNAs were cloned into SpCas9-hGem-T2A-Puromycin/gRNA vector, which is a modified version of SpCas9-T2A-Puromycin/gRNA vector (px459;^116^, Addgene plasmid #62988) similar to that described above. To construct the homology donors plasmids, gBlocks (IDT, Coralville, IA, USA) were synthesized containing either the HALO or eGFP coding sequence flanked by homology arms with ∼200-400 bp homology upstream and downstream of the gRNA target sequence at the *Dppa3* or *Uhrf1* locus, respectively, and then cloned into the NruI site of pUC57-GentR via cut-ligation. ESCs were transfected with equimolar amounts of gRNA and homology donor vectors. Two days after transfection, cells were plated at clonal density and subjected to a transient puromycin selection (1 ug/mL) for 40 h. After 5-6 days, ESCs positive for HALO or eGFP integration were isolated via fluorescence-activated cell sorting (FACS) and plated again at clonal density in ESC media. After 4-5 days, colonies were picked and plated on Optical bottom µClear 96-well plates and re-screened for the correct expression and localization of eGFP or HALO using live-cell spinning-disk confocal imaging. Cells were subsequently genotyped using the aforementioned cell lysis strategy and further validated by Sanger sequencing^113^.

To generate *Dppa3* knockout cells, the targeting strategy entailed the use of two gRNAs with target sites flanking the *Dppa3* locus to excise the entire locus on both alleles. gRNA oligos were cloned into the SpCas9-T2A-PuroR/gRNA vector via cut-ligation. ESCs were transfected with an equimolar amount of each gRNA vector. Two days after transfection, cells were plated at clonal density and subjected to a transient puromycin selection (1 ug/mL) for 40 h. Colonies were picked 6 days after transfection. The triple PCR strategy used for screening is depicted in Supplementary Fig. 3a. Briefly, PCR primers 1F and 4R were used to identify clones in which the *Dppa3* locus had been removed, resulting in the appearance of a ∼350 bp amplicon. To identify whether the *Dppa3* locus had been removed from both alleles, PCRs were performed with primers 1F and 2R or 3F and 4R to amplify upstream or downstream ends of the *Dppa3* locus, which would only be left intact in the event of mono-allelic locus excision. Removal of the *Dppa3* locus was confirmed with Sanger sequencing and loss of *Dppa3* expression was assessed by qRT-PCR.

For CRISPR/Cas gene editing, all transfections were performed using Lipofectamine 3000 (Thermo Fisher Scientific) according to the manufacturer’s instructions. All DNA oligos used for gene editing and screening are listed in Supplementary Table 5.

### Bxb1-mediated recombination and Sleeping Beauty Transposition

To generate stable mESC lines carrying doxycycline inducible forms of *Dppa3* or *Dppa3-mScarlet*, mES cells were first transfected with equimolar amounts of the pSBtet-3xFLAG-Dppa3-IRES-DsRed-PuroR or pSBtet-3xFLAG-Dppa3-mScarlet-PuroR and the Sleeping Beauty transposase, pCMV(CAT)T7-SB100^118^ (Addgene plasmid #34879) vector using Lipofectamine 3000 (Thermo Fisher Scientific) according to manufacturer’s instructions. Two days after transfection, cells were plated at clonal density and subjected to puromycin selection (1 ug/mL) for 5-6 days. To ensure comparable levels of *Dppa3* induction, cells were first treated for 18 h with doxycycline (1 µg/mL) and then sorted with FACS based on thresholded levels of DsRed or mScarlet expression, the fluorescent readouts of successful induction. Post sorting, cells were plated back into media without doxycycline for 7 days before commencing experiments.

To generate stable doxycycline-inducible *Dppa3* hESC lines, hES cells were first transfected with equimolar amounts of the pSBtet-3xFLAG-Dppa3-IRES-DsRed-PuroR and Sleeping Beauty transposase pCMV(CAT)T7-SB100^118^ (Addgene plasmid #34879) vector using using the P3 Primary Cell 4D-NucleofectorTM Kit (V4XP-3012 Lonza) and the 4D-Nucleofector™ Platform (Lonza), program CB-156. Two days after nucleofection, cells were subjected to puromycin selection (1 ug/mL) for subsequent two days, followed by an outgrowth phase of 4days. At this stage, cells were sorted with FACS based on thresholded levels of DsRed expression to obtain two bulk populations of positive stable hESC lines with inducible *Dppa3*.

For the generation of the *Uhrf1*^*GFP/GFP*^ cell line, we used our previously described ESC line with a C-terminal MIN-tag (*Uhrf1*^*attP/attP*^; Bxb1 *attP* site) and inserted the GFP coding sequence as described previously^113^. Briefly, attB-GFP-Stop-PolyA (Addgene plasmid #65526) was inserted into the C-terminal of the endogenous *Uhrf1*^*attP/attP*^ locus by transfection with equimolar amounts of Bxb1 and attB-GFP-Stop-PolyA construct, followed by collection of GFP-positive cells with FACS after 6 days.

### Cellular fractionation and Western Blot

Western blot for T1CM ESCs were performed as described previously^113^ using monoclonal antibody rat anti-TET1 5D6 (1:10)^119^ and polyclonal rabbit anti-H3 (1:5,000; ab1791, Abcam) as loading control. Blots were probed with secondary antibodies goat anti-rat (1:5,000; 112-035-068, Jackson ImmunoResearch) and goat anti-rabbit (1:5,000; 170–6515, Bio-Rad) conjugated to horseradish peroxidase (HRP) and visualized using an ECL detection kit (Thermo Scientific Pierce). Cell fractionation was performed as described previously with minor modifications^120^. 1×10^7^ ESCs were resuspended in 250 µL of buffer A (10 mM HEPES pH 7.9, 10 mM KCl, 1.5 mM MgCl_2_, 0.34 M sucrose, 10% glycerol, 0.1% Triton X-100, 1 mM DTT, 1 mM phenylmethylsulfonyl fluoride (PMSF), 1x mammalian protease inhibitor cocktail (PI; Roche)) and incubated for 5 min on ice. Nuclei were collected by centrifugation (4 min, 1,300 × g, 4 °C) and the cytoplasmic fraction (supernatant) was cleared again by centrifugation (15 min, 20,000 × g, 4 °C). Nuclei were washed once with buffer A, and then lysed in buffer B (3 mM EDTA, 0.2 mM EGTA, 1 mM DTT, 1 mM PMSF, 1x PI). Insoluble chromatin was collected by centrifugation (4 min, 1,700 × g, 4 °C) and washed once with buffer B. Chromatin fraction was lysed with 1x Laemmli buffer and boiled (10 min, 95 °C).

As markers of cytoplasmic and chromatin fractions, alpha-tubulin and histone H3 were detected using monoclonal antibody (mouse anti-alpha-Tubulin, Sigma T9026 or rat anti-Tubulin, Abcam ab6160) and polyclonal antibody (rabbit anti-H3, Abcam ab1791). UHRF1 was visualized by rabbit anti-UHRF1 antibody^75^. Western blots for DNMT1 were performed as described previously using a monoclonal antibody (rat anti-DNMT1, 14F6) or a polyclonal antibody (rabbit anti-DNMT1, Abcam ab87654)^113^. GFP and FLAG tagged proteins were visualized by mouse anti-GFP (Roche) and anti-FLAG M2 antibodies (Sigma, F3165), respectively.

### Quantitative real-time PCR (qRT-PCR) Analysis

Total RNA was isolated using the NucleoSpin Triprep Kit (Macherey-Nagel) according to the manufacturer’s instructions. cDNA synthesis was performed with the High-Capacity cDNA Reverse Transcription Kit (with RNase Inhibitor; Applied Biosystems) using 500 ng of total RNA as input. qRT-PCR assays with oligonucleotides listed in Supplementary Table 5 were performed in 8 µL reactions with 1.5 ng of cDNA used as input. FastStart Universal SYBR Green Master Mix (Roche) was used for SYBR green detection. The reactions were run on a LightCycler480 (Roche).

### LC-MS/MS analysis of DNA samples

Isolation of genomic DNA was performed according to earlier published work^43^. 1.0–5 μg of genomic DNA in 35 μL H_2_O were digested as follows: An aqueous solution (7.5 μL) of 480 μM ZnSO_4_, containing 18.4 U nuclease S1 (Aspergillus oryzae, Sigma-Aldrich), 5 U Antarctic phosphatase (New England BioLabs) and labeled internal standards were added ([^15^N_2_]-cadC 0.04301 pmol, [^15^N_2_,D_2_]-hmdC 7.7 pmol, [D_3_]-mdC 51.0 pmol, [^15^N_5_]-8-oxo-dG 0.109 pmol, [^15^N_2_]-fdC 0.04557 pmol) and the mixture was incubated at 37 °C for 3 h. After addition of 7.5 μl of a 520 μM [Na]_2_-EDTA solution, containing 0.2 U snake venom phosphodiesterase I (Crotalus adamanteus, USB corporation), the sample was incubated for 3 h at 37 °C and then stored at −20 °C. Prior to LC/MS/MS analysis, samples were filtered by using an AcroPrep Advance 96 filter plate 0.2 μm Supor (Pall Life Sciences).

Quantitative UHPLC-MS/MS analysis of digested DNA samples was performed using an Agilent 1290 UHPLC system equipped with a UV detector and an Agilent 6490 triple quadrupole mass spectrometer. Natural nucleosides were quantified with the stable isotope dilution technique. An improved method, based on earlier published work ^43,121^ was developed, which allowed the concurrent analysis of all nucleosides in one single analytical run. The source-dependent parameters were as follows: gas temperature 80 °C, gas flow 15 L/min (N_2_), nebulizer 30 psi, sheath gas heater 275 °C, sheath gas flow 15 L/min (N_2_), capillary voltage 2,500 V in the positive ion mode, capillary voltage −2,250 V in the negative ion mode and nozzle voltage 500 V. The fragmentor voltage was 380 V/ 250 V. Delta EMV was set to 500 V for the positive mode. Chromatography was performed by a Poroshell 120 SB-C8 column (Agilent, 2.7 μm, 2.1 mm × 150 mm) at 35 °C using a gradient of water and MeCN, each containing 0.0085% (v/v) formic acid, at a flow rate of 0.35 mL/min: 0 →4 min; 0 →3.5% (v/v) MeCN; 4 →6.9 min; 3.5 →5% MeCN; 6.9 →7.2 min; 5 →80% MeCN; 7.2 →10.5 min; 80% MeCN; 10.5 →11.3 min; 80 →0% MeCN; 11.3 →14 min; 0% MeCN. The effluent up to 1.5 min and after 9 min was diverted to waste by a Valco valve. The autosampler was cooled to 4 °C. The injection volume was amounted to 39 μL. Data were processed according to earlier published work^43^.

### RNA-seq and Reduced Representation Bisulfite Sequencing (RRBS)

For RNA-seq, RNA was isolated using the NucleoSpin Triprep Kit (Machery-Nagel) according to the manufacturer’s instructions. Digital gene expression libraries for RNA-seq were produced using a modified version of single-cell RNA barcoding sequencing (SCRB-seq) optimized to accommodate bulk cells^122^ in which a total of 70 ng of input RNA was used for the reverse-transcription of individual samples. For RRBS, genomic DNA was isolated using the QIAamp DNA Mini Kit (QIAGEN), after an overnight lysis and proteinase K treatment. RRBS library preparation was performed as described previously^123^, with the following modifications: bisulfite treatment was performed using the EZ DNA Methylation-Gold™ Kit (Zymo Research Corporation) according to the manufacturer’s protocol except libraries were eluted in 2 x 20 µL M-elution buffer. RNA-seq and RRBS libraries were sequenced on an Illumina HiSeq 1500.

### Targeted Bisulfite Amplicon (TaBA) Sequencing

Genomic DNA was isolated from 10^6^ cells using the PureLink Genomic DNA Mini Kit (Thermo Fisher Scientific) according to the manufacturer’s instructions. The EZ DNA Methylation-Gold Kit (Zymo Research) was used for bisulfite conversion according to the manufacturer’s instructions with 500 ng of genomic DNA used as input and the modification that bisulfite converted DNA was eluted in 2 x 20 µL Elution Buffer.

The sequences of the locus specific primers (Supplementary Table 5) were appended with Illumina TruSeq and Nextera compatible overhangs. The amplification of bisulfite converted DNA was performed in 25 µL PCR reaction volumes containing 0.4 µM each of forward and reverse primers, 2 mM Betaiinitialne (Sigma-Aldrich, B0300-1VL), 10 mM Tetramethylammonium chloride solution (Sigma-Aldrich T3411-500ML), 1x MyTaq Reaction Buffer, 0.5 units of MyTaq HS (Bioline, BIO-21112), and 1 µL of the eluted bisulfite converted DNA (∼12.5 ng). The following cycling parameters were used: 5 min for 95 °C for initial denaturation and activation of the polymerase, 40 cycles (95 °C for 20 s, 58 °C for 30 s, 72 °C for 25 s) and a final elongation at 72 °C for 3 min. Agarose gel electrophoresis was used to determine the quality and yield of the PCR.

For purifying amplicon DNA, PCR reactions were incubated with 1.8x volume of CleanPCR beads (CleanNA, CPCR-0005) for 10 min. Beads were immobilized on a DynaMag™-96 Side Magnet (Thermo Fisher, 12331D) for 5 min, the supernatant was removed, and the beads washed 2x with 150 µL 70% ethanol. After air drying the beads for 5 min, DNA was eluted in 15 µL of 10 mM Tris-HCl pH 8.0. Amplicon DNA concentration was determined using the Quant-iT™ PicoGreen™ dsDNA Assay Kit (Thermo Fisher, P7589) and then diluted to 0.7 ng/µL. Thereafter, indexing PCRs were performed in 25 µL PCR reaction volumes containing 0.08 µM (1 µL of a 2 µM stock) each of i5 and i7 Indexing Primers, 1x MyTaq Reaction Buffer, 0.5 units of MyTaq HS (Bioline, BIO-21112), and 1 µL of the purified PCR product from the previous step. The following cycling parameters were used: 5 min for 95 °C for initial denaturation and activation of the polymerase, 40 cycles (95 °C for 10 s, 55 °C for 30 s, 72 °C for 40 s) and a final elongation at 72 °C for 5 min. Agarose gel electrophoresis was used to determine the quality and yield of the PCR. An aliquot from each indexing reaction (5 µL of each reaction) was then pooled and purified with CleanPCR magnetic beads as described above and eluted in 1 µL x Number of pooled reactions. Concentration of the final library was determined using PicoGreen and the quality and size distribution of the library was assessed with a Bioanalyzer. Dual indexed TaBA-seq libraries were sequenced on an Illumina MiSeq in 2×300 bp output mode.

### RNA-seq processing and analysis

RNA-seq libraries were processed and mapped to the mouse genome (mm10) using the zUMIs pipeline^124^. UMI count tables were filtered for low counts using HTSFilter^125^. Differential expression analysis was performed in R using DESeq2^126^ and genes with an adjusted P < 0.05 were considered to be differentially expressed. Hierarchical clustering was performed on genes differentially expressed in TET mutant ESCs respectively, using k-means clustering (k = 4) in combination with the ComplexHeatmap R-package^127^. Principal component analysis was restricted to genes differentially expressed during wild-type differentiation and performed using all replicates of wild-type, TET mutant, and Dppa3KO ESCs.

### RRBS alignment and analysis

Raw RRBS reads were first trimmed using Trim Galore (v.0.3.1) with the ‘--rrbs’ parameter. Alignments were carried out to the mouse genome (mm10) using bsmap (v.2.90) using the parameters ‘-s 12 -v 10 -r 2 -I 1’. CpG-methylation calls were extracted from the mapping output using bsmaps methratio.py. Analysis was restricted to CpG with a coverage > 10. methylKit ^128^ was used to identify differentially methylated regions between the respective contrasts for the following genomic features: 1) all 1-kb tiles (containing a minimum of three CpGs) detected by RRBS; 2) Repeats (defined by Repbase); 3) gene promoters (defined as gene start sites −2kb/+2kb); and 4) gene bodies (defined as longest isoform per gene) and CpG islands (as defined by ^129^). Differentially methylated regions were identified as regions with P < 0.05 and a difference in methylation means between two groups greater than 20%. Principal component analysis of global DNA methylation profiles was performed on single CpGs using all replicates of wild-type, T1KO and T1CM ESCs and EpiLCs.

### Chromatin immunoprecipitation (ChIP) and Hydroxymethylated-DNA immunoprecipitation (hMeDIP) alignment and analysis

ChIP–seq reads for TET1 binding in ESCs and EpiLCs were downloaded from GSE57700^45^ and PRJEB19897^44^, respectively. hMeDIP reads for wild-type ESCs and T1KO ESCs were download from PRJEB13096^44^. Reads were aligned to the mouse genome (mm10) with Bowtie (v.1.2.2) with parameters ‘-a -m 3 -n 3 --best --strata’. Subsequent ChIP–seq analysis was carried out on data of merged replicates. Peak calling and signal pile up was performed using MACS2 callpeak^130^ with the parameters ‘--extsize 150’ for ChIP, ‘--extsize 220’ for hMeDIP, and ‘--nomodel -B --nolambda’ for all samples. Tag densities for promoters and 1kb Tiles were calculated using the deepTools2 computeMatrix module^131^. TET1 bound genes were defined by harboring a TET1 peak in the promoter region (defined as gene start sites −2kb/+2kb).

### Immunofluorescence staining

For immunostaining, naïve ESCs were grown on coverslips coated with Geltrex (Life Technologies) diluted 1:100 in DMEM/F12 (Life Technologies), thereby allowing better visualization of the cytoplasm during microscopic analysis. All steps during immunostaining were performed at room temperature. Coverslips were rinsed two times with PBS (pH 7.4; 140 mM NaCl, 2.7 mM KCl, 6.5 mM Na_2_HPO_4_, 1.5 mM KH_2_PO_4_) prewarmed to 37 °C, cells fixed for 10 min with 4% paraformaldehyde (pH 7.0; prepared from paraformaldehyde powder (Merck) by heating in PBS up to 60 °C; store at -20 °C), washed three times for 10 min with PBST (PBS, 0.01% Tween20), permeabilized for 5 min in PBS supplemented with 0.5% Triton X-100, and washed two times for 10 min with PBS. Primary and secondary antibodies were diluted in blocking solution (PBST, 4% BSA). Coverslips were incubated with primary and secondary antibody solutions in dark humid chambers for 1 h and washed three times for 10 min with PBST after primary and secondary antibodies. For DNA counterstaining, coverslips were incubated 6 min in PBST containing a final concentration of 2 µg/mL DAPI (Sigma-Aldrich) and washed three times for 10 min with PBST. Coverslips were mounted in antifade medium (Vectashield, Vector Laboratories) and sealed with colorless nail polish.

Following primary antibodies were used: polyclonal rabbit anti-DPPA3 (1:200; ab19878, Abcam) and monoclonal rabbit anti-UHRF1 (1:250; D6G8E, Cell Signaling). Following secondary antibodies were used: polyclonal donkey anti-rabbit conjugated to Alexa 488 (1:500; 711-547-003, Dianova), polyclonal donkey anti-rabbit conjugated to Alexa 555 (1:500; A31572, Invitrogen), and polyclonal donkey anti-rabbit conjugated to DyLight fluorophore 594 (1:500; 711-516-152, Dianova).

### Immunofluorescence and Live-cell imaging

For immunofluorescence, stacks of optical sections were collected on a Nikon TiE microscope equipped with a Yokogawa CSU-W1 spinning disk confocal unit (50 μm pinhole size), an Andor Borealis illumination unit, Andor ALC600 laser beam combiner (405nm/488nm/561nm/640nm), Andor IXON 888 Ultra EMCCD camera, and a Nikon 100x/1.45 NA oil immersion objective. The microscope was controlled by software from Nikon (NIS Elements, ver. 5.02.00). DAPI or fluorophores were excited with 405 nm, 488 nm, or 561 nm laser lines and bright-field images acquired using Nikon differential interference contrast optics. Confocal image z-stacks were recorded with a step size of 200 nm, 16-bit image depth, 1 × 1 binning, a frame size of 1024 × 1024 pixels, and a pixel size of 130 nm. Within each experiment, cells were imaged using the same settings on the microscope (camera exposure time, laser power, and gain) to compare signal intensities between cell lines. For live-cell imaging, cells were plated on Geltrex-coated glass bottom 2-well imaging slides (Ibidi). Both still and timelapse images were acquired on the Nikon spinning disk system described above equipped with an environmental chamber maintained at 37 °C with 5% CO_2_ (Oko Labs), using a Nikon 100x/1.45 NA oil immersion objective and a Perfect Focus System (Nikon). Images were acquired with the 488, 561, and 640 nm laser lines, full-frame (1024 × 1024) with 1 × 1 binning, and with a pixel size of 130 nm. Transfection of a RFP-PCNA vector^132^ was used to identify cells in S-phase. For DNA staining in live cells, cells were exposed to media containing 200 nM SiR-DNA (Spirochrome) for at least 1 h before imaging. For imaging endogenous DPPA3-HALO in live cells, cells were treated with media containing 50 nM HaloTag-TMR fluorescent ligand (Promega) for 1 h. After incubation, cells were washed 3x with PBS before adding back normal media. Nuclear export inhibition was carried out using media containing 20 nM leptomycin-B (Sigma-Aldrich).

### Image analysis

For immunofluorescence images, Fiji software (ImageJ 1.51j)^133,134^ was used to analyze images and create RGB stacks. For analysis of live-cell imaging data, CellProfiler Software (version 3.0)^135^ was used to quantify fluorescence intensity in cells stained with SiR-DNA. CellProfiler pipelines used in this study are available upon request. In brief, the SiR-DNA signal was used to segment ESC nuclei. Mean fluorescence intensity of GFP was measured both inside the segmented area (nucleus) and in the area extending 4-5 pixels beyond the segmented nucleus (cytoplasm). GFP fluorescence intensity was normalized by subtracting the experimentally-determined mean background intensity and background-subtracted GFP intensities were then used for all subsequent quantifications shown in Fig. 4 and Supplementary Figs. 4h, 5h, and 6b,c.

### Fluorescence Recovery After Photobleaching (FRAP)

For FRAP assays, cells cultivated on Geltrex-coated glass bottom 2-well imaging slides (Ibidi) were imaged in an environmental chamber maintained at 37 °C with 5% CO_2_ either using the Nikon system mentioned above equipped with a FRAPPA photobleaching module (Andor) or on an Ultraview-Vox spinning disk system (Perkin-Elmer) including a FRAP Photokinesis device mounted to an inverted Axio Observer D1 microscope (Zeiss) equipped with an EMCCD camera (Hamamatsu) and a 63x/1.4 NA oil immersion objective, as well as 405, 488 and 561 nm laser lines.

For endogenous UHRF1-GFP FRAP, eight pre-bleach images were acquired with the 488 nm laser, after which an area of 4 x 4 pixels was irradiated for a total of 16 ms with a focused 488 nm laser (leading to a bleached spot of ∼1 μm) to bleach a fraction of GFP-tagged molecules within cells, and then recovery images were acquired every 250 ms for 1-2 min. Recovery analysis was performed in Fiji. Briefly, fluorescence intensity at the bleached spot was measured in background-subtracted images, then normalized to pre-bleach intensity of the bleached spot, and normalized again to the total nuclear intensity in order to account for acquisition photobleaching. Images of cells with visible drift were discarded.

### *Xenopus* egg extracts

The interphase extracts (low-speed supernatants (LSS)) were prepared as described previously^66^. After thawing, LSS were supplemented with an energy regeneration system (5 μg/ml creatine kinase, 20 mM creatine phosphate, 2 mM ATP) and incubated with sperm nuclei at 3,000-4,000 nuclei per μl. Extracts were diluted 5-fold with ice-cold CPB (50 mM KCl, 2.5 mM MgCl2, 20 mM HEPES-KOH, pH 7.7) containing 2% sucrose, 0.1% NP-40 and 2 mM NEM, overlaid onto a 30% sucrose/CPB cushion, and centrifuged at 15,000g for 10min. The chromatin pellet was resuspended in SDS sample buffer and analyzed by SDS-PAGE. GST-mDPPA3 was added to egg extracts at 50 ng/μl at final concentration.

### Monitoring DNA methylation in *Xenopus* egg extracts

DNA methylation was monitored by the incorporation of S-[methyl-3H]-adenosyl-L-methionine, incubated at room temperature, and the reaction was stopped by the addition of CPB containing 2% sucrose up to 300 μl. Genomic DNA was purified using a Wizard Genomic DNA purification kit (Promega) according to the manufacturer’s instructions. Incorporation of radioactivity was quantified by liquid synchillation counter.

### Plasmid construction for Recombinant mDPPA3

To generate GST-tagged mDPPA3 expression plasmids, mDPPA3 fragment corresponding to full-length protein was amplified by PCR using mouse DPPA3 cDNA and primers 5’-TTAGCAGCCGGATCCCTAATTCTTCCCGATTTTCGCA-3’ and 5’-CGTGGATCCCCGAATTCCATGGAGGAACCATCAGAGAAAGTC’. The resulting DNA fragment was cloned into pGEX4T-3 vector digested with EcoRI and SalI using an In-Fusion HD Cloning Kit.

### Protein expression and purification

For protein expression in Escherichia coli (BL21-CodonPlus), the mDPPA3 genes were transferred to pGEX4T-3 vector as described above. Protein expression was induced by the addition of 0.1 mM Isopropyl β–D-1-thiogalactopyranoside (IPTG) to media followed by incubation for 12 hour at 20°C. For purification of Glutathione S transferase (GST) tagged proteins, cells were collected and re-suspended in Lysis buffer (20 mM HEPES-KOH (pH 7.6), 0.5M NaCl, 0.5 mM EDTA, 10% glycerol, 1 mM DTT) supplemented with 0.5% NP40 and protease inhibitors, and were then disrupted by sonication on ice. After centrifugation, the supernatant was applied to Glutathione Sepharose (GSH) beads (GE Healthcare) and rotated at 4 °C for 2 hour. Beads were then washed three times with Wash buffer 1 (20 mM Tris-HCl (pH 8.0), 150 mM NaCl, 1% TritionX-100, 1 mM DTT) three times and with Wash buffer 2 (100 mM Tris-HCl (pH 7.5), 100 mM NaCl) once. Bound proteins were eluted in Elution buffer (100 mM Tris-HCl (pH 7.5), 100 mM NaCl, 5% glycerol, 1 mM DTT) containing 42 mM reduced Glutathione and purified protein was loaded on PD10 desalting column equilibrated with EB buffer (10 mM HEPES/KOH at pH 7.7, 100 mM KCl, 0.1 mM CaCl2, 1 mM MgCl2) containing 1 mM DTT, and then concentrated by Vivaspin (Millipore).

### Data collection for the presence of TET1, UHRF1, DNMT1 and DPPA3 throughout metazoa

Megabat (*Pteropus vampyrus*) protein sequences for TET1 (ENSVPAP00000010999), UHRF1 (ENSPVAP00000002809), DNMT1 (ENSPVAP00000003477) and DPPA3 (ENSVPAP00000004109) were subjected to three iterations of PSI-blast ^136^, respectively. Subsequently, result tables were filtered to contain correct gene names using a custom python script. Presence of the proteins throughout metazoa was visualized using iTOL^137^.

### Chromatin immunoprecipitation coupled to Mass Spectrometry and Proteomics data analysis

For Chromatin immunoprecipitation coupled to Mass Spectrometry (ChIP-MS), whole cell lysates of the doxycycline inducible *Dppa3*-FLAG mES cells were used by performing three separate immunoprecipitations with an anti-FLAG antibody and three samples with a control IgG. Proteins were digested on the beads after the pulldown and desalted subsequently on StageTips with three layers of C18^138^. Here, peptides were separated by liquid chromatography on an Easy-nLC 1200 (Thermo Fisher Scientific) on in-house packed 50 cm columns of ReproSil-Pur C18-AQ 1.9-µm resin (Dr. Maisch GmbH). Peptides were then eluted successively in an ACN gradient for 120 min at a flow rate of around 300 nL/min and were injected through a nanoelectrospray source into a Q Exactive HF-X Hybrid Quadrupole-Orbitrap Mass Spectrometer (Thermo Fisher Scientific). After measuring triplicates of a certain condition, an additional washing step was scheduled. During the measurements, the column temperature was constantly kept at 60 °C while after each measurement, the column was washed with 95% buffer B and subsequently with buffer A. Real time monitoring of the operational parameters was established by SprayQc^139^ software. Data acquisition was based on a top10 shotgun proteomics method and data-dependent MS/MS scans. Within a range of 400-1650 m/z and a max. injection time of 20 ms, the target value for the full scan MS spectra was 3 × 10^6^ and the resolution at 60,000.

The raw MS data was then analyzed with the MaxQuant software package (version 1.6.0.7)^140^. The underlying FASTA files for peak list searches were derived from Uniprot (UP000000589_10090.fasta and UP000000589_10090 additional.fasta, version June 2015) and an additional modified FASTA file for the FLAG-tagged *Dppa3* in combination with a contaminants database provided by the Andromeda search engine^141^ with 245 entries. During the MaxQuant-based analysis the “Match between runs” option was enabled and the false discovery rate was set to 1% for both peptides (minimum length of 7 amino acids) and proteins. Relative protein amounts were determined by the MaxLFQ algorithm^142^, with a minimum ratio count of two peptides.

For the downstream analysis of the MaxQuant output, the software Perseus^143^ (version 1.6.0.9) was used to perform Student’s *t*-test with a permutation-based FDR of 0.05 and an additional constant S0 = 1 in order to calculate fold enrichments of proteins between triplicate chromatin immunoprecipitations of anti-FLAG antibody and control IgG. The result was visualized in a scatter plot.

For GO analysis of biological processes the Panther classification system was used^143^. For the analysis, 131 interactors of DPPA3 were considered after filtering the whole amount of 303 significant interactors for a p-value of at least 0.0015 and 3 or more identified peptides. The resulting GO groups were additionally filtered for a fold enrichment of observed over expected amounts of proteins of at least 4 and a p-value of 5.30 E-08.

### *Dppa3* overexpression in medaka embryos and immunostaining

Medaka d-rR strain was used. Medaka fish were maintained and raised according to standard protocols. Developmental stages were determined based on a previous study^144^. *Dppa3* and mutant *Dppa3* (R107E) mRNA were synthesized using HiScribe T7 ARCA mRNA kit (NEB, E2060S), and purified using RNeasy mini kit (QIAGEN, 74104). *Dppa3* or mutant *Dppa3* (R107E) mRNA was injected into the one-cell stage (stage 2) medaka embryos. After 7 hours of incubation at 28 °C, the late blastula (stage 11) embryos were fixed with 4% PFA in PBS for 2 hours at room temperature, and then at 4 °C overnight. Embryos were dechorionated, washed with PBS, and permeabilized with 0.5% Triton X-100 in PBS for 30 minutes at room temperature. DNA was denatured in 4 M HCl for 15 minutes at room temperature, followed by neutralization in 100 mM Tris-HCl (pH 8.0) for 20 minutes. After washing with PBS, embryos were blocked in blocking solution (2% BSA, 1%DMSO, 0.2% Triton X-100 in PBS) for 1 hour at room temperature, and then incubated with 5-methylcytosine antibody (1:200; Active Motif #39649) at 4 °C overnight. The embryos were washed with PBSDT (1% DMSO, 0.1% Triton X-100 in PBS), blocked in blocking solution for 1 hours at room temperature, and incubated with Alexa Fluor 555 goat anti-mouse 2nd antibody (1:500; ThermoFisher Scientific #A21422) at 4 °C overnight. After washing with PBSDT, cells were mounted on slides and examined under a fluorescence microscope.

### Fluorescence Three Hybrid (F3H) assay

The F3H assay was performed as described previously^82^. In brief, BHK cells containing multiple lac operator repeats were transiently transfected with the respective GFP- and dsRed-constructs on coverslips using PEI and fixed with 3.0% formaldehyde 24 hrs after transfection. For DNA counterstaining, coverslips were incubated in a solution of DAPI (200 ng/ml) in PBS-T and mounted in Vectashield. Images were collected using a Leica TCS SP5 confocal microscope. To quantify the interactions within the lac spot, the following intensity ratio was calculated for each cell in order to account for different expression levels: (mCherry_spot_–mCherry_background_)/(GFP_spot_–GFP_background_).

### Microscale Thermophoresis (MST)

For MST measurements, mUHRF1 C-terminally tagged with GFP- and 6xHis-tag was expressed in HEK 293T cells and then purified using Qiagen Ni-NTA beads (Qiagen #30230). Recombinant mDPPA3 WT and 1-60 were purified as described above. Purified UHRF1 (200 nM) was mixed with different concentrations of purified DPPA3 (0.15 nM to 5 µM) followed by a 30 min incubation on ice. The samples were then aspirated into NT.115 Standard Treated Capillaries (NanoTemper Technologies) and placed into the Monolith NT.115 instrument (NanoTemper Technologies). Experiments were conducted with 80% LED and 80% MST power. Obtained fluorescence signals were normalized (F_norm_) and the change in F_norm_ was plotted as a function of the concentration of the titrated binding partner using the MO. Affinity Analysis software version 2.1 (NanoTemper Technologies). For fluorescence normalization (F_norm_ = F_hot_/F_cold_), the manual analysis mode was selected and cursors were set as follows: F_cold_ = -1 to 0 s, F_hot_ = 10 to 15 s. The Kd was obtained by fitting the mean F_norm_ of eight data points (four independent replicates, each measured as a technical duplicate).

### RICS

Data for Raster Image Correlation Spectroscopy (RICS) was acquired on a home-built laser scanning confocal setup equipped with a pulsed interleaved excitation (PIE) system as used elsewhere^145^. Samples were excited using pulsed lasers at 470 (Picoquant) and 561 nm (Toptica Photonics), synchronized to a master clock, and then delayed ∼20ns relative to one another to achieve PIE. Laser excitation was separated from descanned fluorescence emission by a Di01-R405/488/561/635 polychroic mirror (Semrock, AHF Analysentechnik) and eGFP and mScarlet fluorescence emission was separated by a 565 DCXR dichroic mirror (AHF Analysentechnik) and collected on avalanche photodiodes, a Count Blue (Laser Components) and a SPCM-AQR-14 (Perkin-Elmer) with 520/40 and a 630/75 emission filters (Chroma, AHF Analysentechnik). Detected photons were recorded by time-correlated single-photon counting.

The alignment of the system was verified prior to each measurement session by performing FCS with PIE of a mixture of Atto-488 and Atto565 dyes, excited with pulsed 470 and 561 nm lasers set to 10 μW (measured in the collimated space before entering the galvo-scanning mirror system), 1 μm above the surface of the coverslip^146^. Cells were plated on Ibidi two-well glass bottom slides, and induced with doxycycline overnight prior to measurements. Scanning was performed in cells at maintained at 37 °C with a stage top incubator, with a total field of view of 12 × 12 µm, composed of 300 pixels x 300 lines a pixel size of 40 nm, a pixel dwell time of 11 µs, a line time 3.33 ms, at one frame per second, for 100-200 seconds. Pulsed 470 and 561 nm lasers were adjusted to 4 and 5 μW respectively.

Image analysis was done using the Pulsed Interleaved Excitation Analysis with Matlab (PAM) software^147^. Briefly, time gating of the raw photon stream was performed by selecting only photons emitted on the appropriate detector after pulsed excitation, thereby allowing to generate cross-talk free images for each channel. Then, using Microtime Image Analysis (MIA), slow fluctuations were removed by subtracting a moving average of 3 frames, and a region of interest corresponding to the nucleus was selected, excluding nucleoli and dense aggregates. The spatial autocorrelation and cross-correlation functions (SACF and SCCF) were calculated as done previously^148^ using arbitrary region RICS:

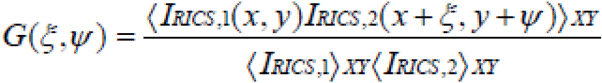

where ξ and ψ are the correlation lags in pixel units along the x- and y-axis scan directions. The correlation function was then fitted to a two-component model (one mobile and one immobile component) in MIAfit:

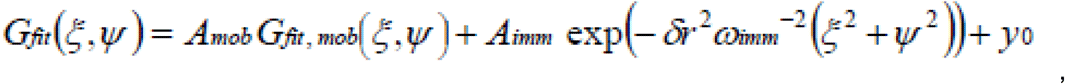

where

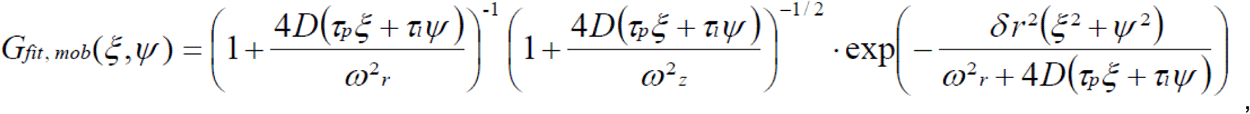

which yields parameters such as the diffusion coefficient (D) and the amplitudes of the mobile and immobile fractions (A_mob_ and A_imm_). The average number of mobile molecules per excitation volume on the RICS timescale was determined by:

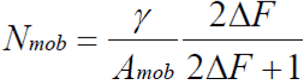

where γ is a factor pertaining to the 3D Gaussian shape of the PSF, and 2ΔF/(2ΔF+1) is a correction factor when using a moving average subtraction prior to calculating the SACF. The bound fraction is the contribution of particles which remain visible during the acquisition of 5-10 lines of the raster scan, corresponding to ∼30 ms. The cross-correlation model was fitted to the cross-correlation function, and the extent of cross-correlation was calculated from the amplitude of the mobile fraction of the cross-correlation fit divided by the amplitude of the mobile fraction of the autocorrelation fit of DPPA3-mScarlet.

### Data and Code Availability

Sequencing data reported in this paper are available at ArrayExpress (EMBL-EBI) under accessions E-MTAB-6785 (wild-type and Tet catalytic mutants RRBS), E-MTAB-6797 (RNA-seq), E-MTAB-6800 (Dppa3KO RRBS). The raw mass spectrometry proteomics data from the FLAG-DPPA3 pulldown will be deposited at the ProteomeXchange Consortium via the PRIDE partner repository. Significant interactors of FLAG-DPPA3 in ESCs can be found in Supplementary Table 3.

### Statistics and reproducibility

No statistical methods were used to predetermine sample size, the experiments were not randomized, and the investigators were not blinded to allocation during experiments and outcome assessment. Blinding was not implemented in this study as analysis was inherently objective in the overwhelming majority of experiments. For microscopy analysis, where possible, experimenter bias was avoided by selecting fields of view (or individual cells) for acquisition of UHRF1-GFP or DNMT1-GFP signal using the DNA stain (or another marker not being directly assessed in the experiment e.g. DsRed/mScarlet as a readout of Dppa3 induction or RFP-PCNA). To further reduce bias, imaging analysis was subsequently performed indiscriminately on all acquired images using semi-automated analysis pipelines (either with CellProfiler or Fiji scripts). All the experimental findings were reliably reproduced in independent experiments as indicated in the Figure legends. The number of replicates used in each experiment are described in the Figure legends and/or in the Methods section, as are the Statistical tests used. P values values or adjusted P values are given where possible. Unless otherwise indicated, all statistical calculations were performed using R Studio 1.2.1335. Next-generation sequencing experiments include at least two independent biological replicates. RNA-seq experiments include *n*= 4 biological replicates comprised of *n*=2 independently cultured samples from 2 clones (for T1CM, T2CM, T12CM ESCs and EpiLCs) or 4 independently cultured samples (for wild-type ESCs and EpiLCs). For RRBS experiments, data are derived from *n*= 2 biological replicates. For bisulfite sequencing of LINE-1 elements *n*= 2 biological replicates were analyzed from 2 independent clones for T1CM, T2CM, T12CM, and Dppa3KO ESCs or 2 independent cultures for wt ESCs. LC-MS/MS quantification was performed on at least 4 biological replicates comprising at least 2 independently cultured samples (usually even more) from *n* = 2 independent clones (T1CM, T2CM, T12CM, and Dppa3KO ESCs) or 4 independently cultured samples (wild-type ESCs and cell lines shown in Fig. 5d).

## Acknowledgements

We would like to thank Dr. H. Blum, Dr. S. Krebs (Laboratory for Functional Genome Analysis, LMU Munich), and Dr. A. Brachmann (Faculty of Biology, Department of Genetics, LMU Munich) for next generation sequencing services. We further thank Dr. Ralf Heermann (Bioanalytics core facility, LMU Munich) for help with the MST measurements. We thank V. Laban, C. Kirschner, T.J. Fischer, D. Nixdorf and V.B. Ozan for help with experiments, J. Koch for technical assistance, Dr. M. Mann and Dr. I. Paron for mass spectrometry and Dr. I. Hellmann for advice on data analysis and providing computational infrastructure. We would like to thank Dr. S. Stricker and Dr. I. Solovei for helpful discussions and constructive criticism on the manuscript and E. Ntouliou for critical reading of the manuscript. We thank Dr. Feng Zhang for providing the pSpCas9(BB)-2A-Puro (PX459) and SpCas9-T2A-GFP (PX458) (Addgene plasmids #62988 and #48138) plasmids, Dr. Eric Kowarz for the gift of the pSB-tetPur (Addgene plasmid #60507) construct, and Dr. Zsuzsanna Izsvak for the gift of the pCMV(CAT)T7-SB100 (Addgene plasmid #34879). J.R. and M.D.B. are fellows of the International Max Planck Research School for Molecular Life Sciences (IMPRS-LS). C.B.M gratefully acknowledges the support of the Fulbright Commission and the late Dr. Glenn Cuomo. F.R.T. thanks the Boehringer Ingelheim Fonds and J.R. the Fonds de Recherche du Québec en Santé for a PhD fellowship.

## Funding

The work was funded by the Deutsche Forschungsgemeinschaft (DFG grants SFB1064/A22 to S.B, SFB1064/A17 and SFB1243/A01 to H.L., and SFB1243/A14 to W.E.).

## Authors Contributions

C.B.M. and S.B. designed and conceived the study. S.B. and H.L. supervised the study. C.M., S.B., and H.L. prepared the manuscript with the help of M.D.B. C.B.M. performed cellular and molecular experiments. C.B.M. generated cell lines with help from M.Y.. C.B.M. performed RRBS and RNA-Seq with help and supervision from C.Z., S.B., and W.E.. J.R. and C.B.M. performed live-cell microscopy and photobleaching analyses. I.G., J.R. and C.B.M. performed RICS experiments under the supervision of D.L.. C.T. performed MST and F3H assays. A.N. performed *Xenopus* experiments under the supervision of M.N.. R.N. performed the experiments in medaka embryos under the supervision of H.T.. M.D.B. and P.S. helped with cell line validation and performed fluorescence microscopy analysis. W.Q. performed the biochemical analyses with assistance from A.A. M.M performed hESC experiments. E.U. conducted proteomics experiments and analyses under the guidance of M.W. F.R.T. and E.P. quantified modified cytosines by LC-MS/MS with the supervision by T.C. S.B. performed data analysis. All authors read, discussed, and approved the manuscript.

## Competing interests

The authors declare no competing interests.

## SUPPLEMENTARY FIGURES

### This PDF includes

Supplementary Fig. 1

Supplementary Fig. 2

Supplementary Fig. 3

Supplementary Fig. 4

Supplementary Fig. 5

Supplementary Fig. 6

Supplementary Fig. 7

Supplementary Fig. 8

### Other Supplementary Information for this manuscript not in this PDF

Supplementary Table 1

Supplementary Table 2

Supplementary Table 3

Supplementary Table 4

Supplementary Table 5

**Supplementary Figure 1:**
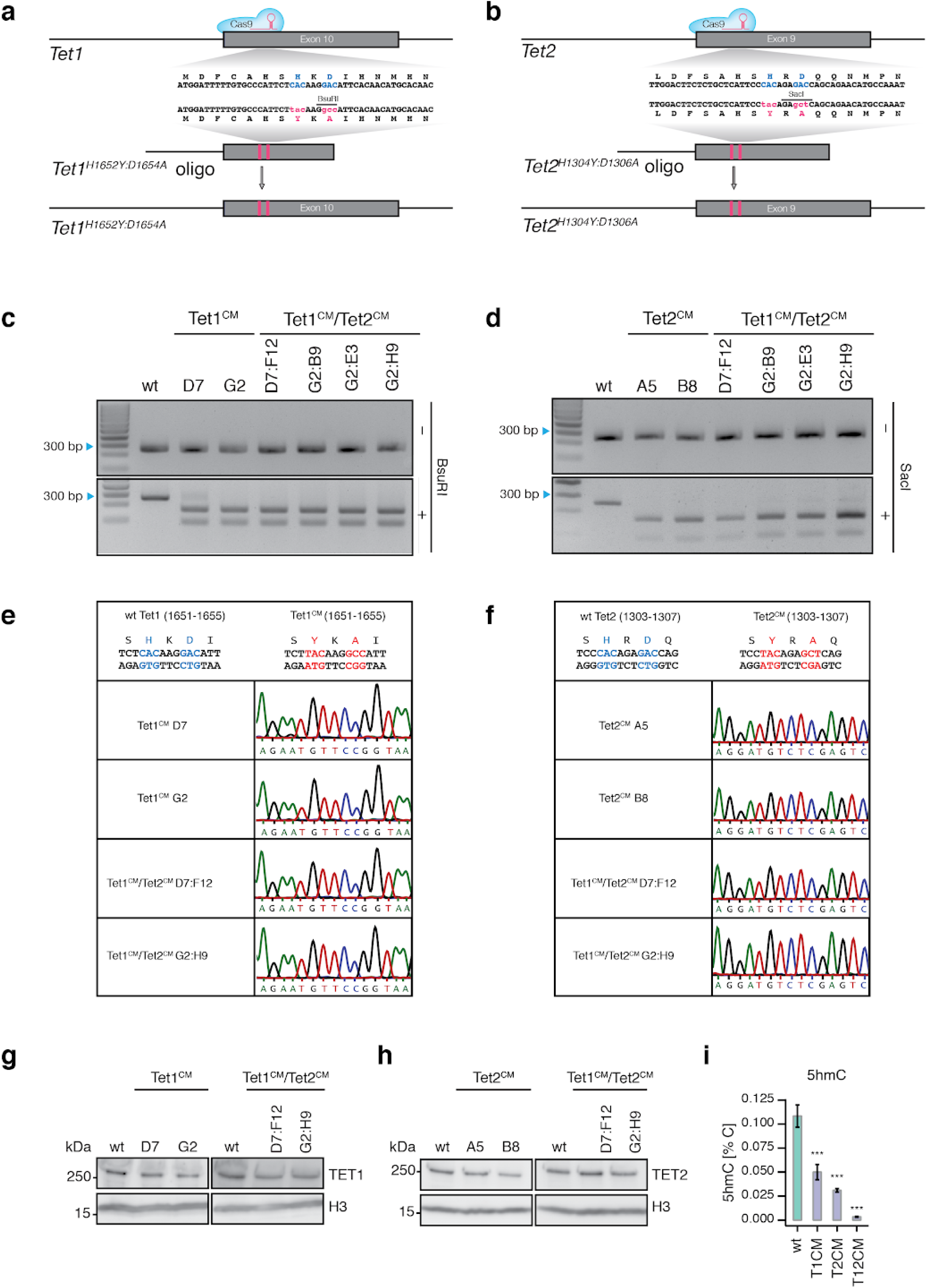
Generation and characterization of T1CM, T2CM, and T12CM mESCs. **a,b**, Schematic representation of CRISPR/Cas9 gene editing strategy used to mutate the catalytic center (HxD) of *Tet1* (**a**) and *Tet2* (**b**). gRNA target sequences and restriction enzyme recognition sites for restriction fragment length polymorphism (RFLP) screening are shown (See Supplementary Table 5). **c,d**, Genotyping using RFLP analysis of the *Tet1* (**c**) and *Tet2* (**d**) locus. **e,f**, Sanger sequencing results confirming the successful insertion of point mutations, YxA, at the *Tet1* (**g**) and *Tet2* (**h**) locus, HxD. **g,h**, Immunoblot detection of endogenous TET1 (**g**) and TET2 (**h**) protein levels in T1CM, T2CM, and T12CM mESCs. **i**, DNA modification levels as percentage of total cytosines in wt (*n* = 24), T1CM (*n* = 8), T2CM (*n* = 12), and T12CM (*n* = 11) mESCs. Depicted are mean values ± standard deviation; *P < 0.05, **P < 0.01, ***P < 0.005 to wt as determined using a one-way ANOVA followed by a post-hoc Tukey HSD test.

**Supplementary Figure 2:**
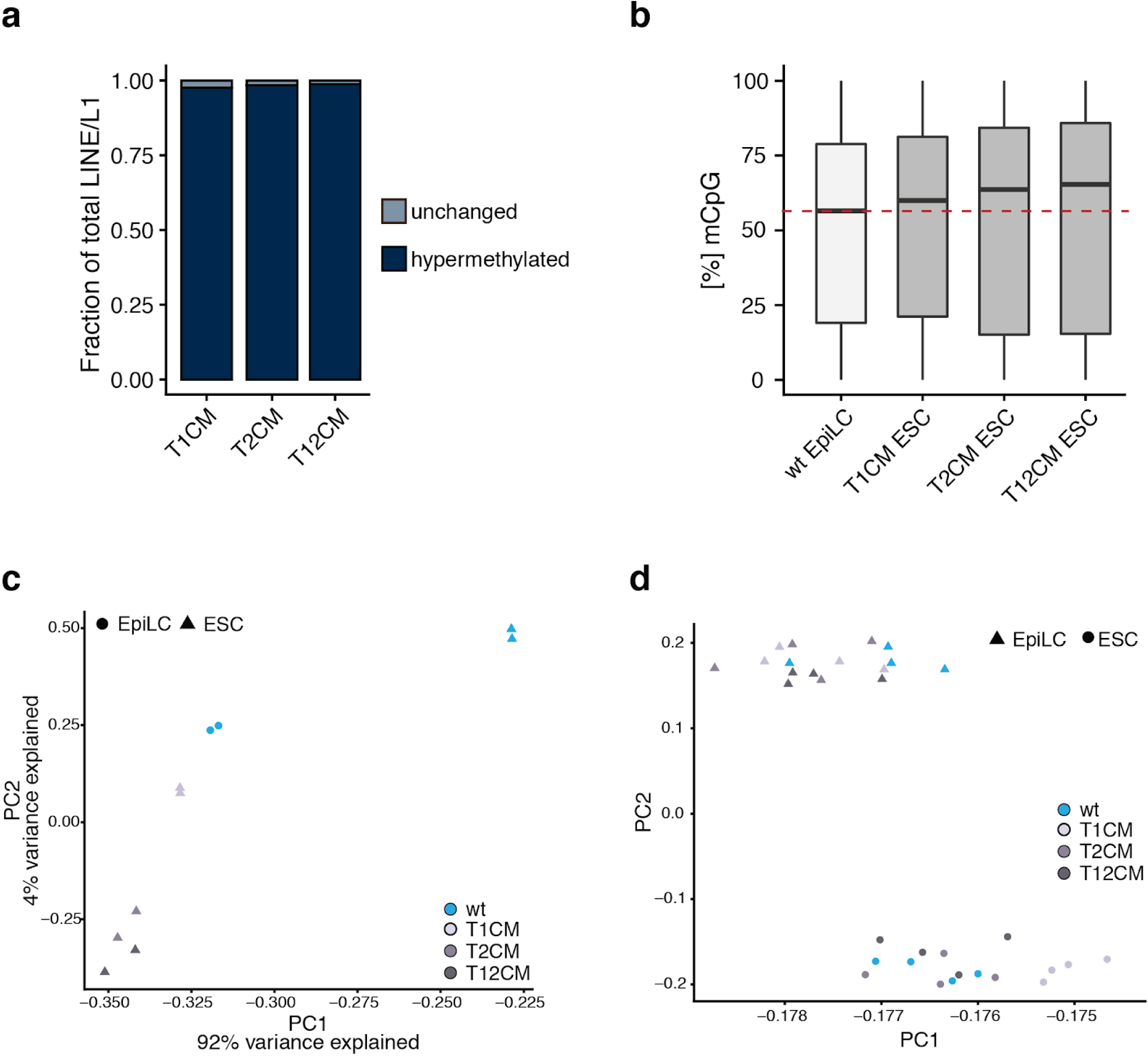
Methylome and transcriptome analysis of T1CM, T2CM, and T12CM ESCs. **a**, Relative proportion of DNA hypermethylation (q value < 0.05; absolute methylation difference > 20%) at LINE1/L1 elements in T1CM, T2CM, and T12CM ESCs compared to wt ESCs. **b**, Loss of TET catalytic activity in ESCs results in similar or higher DNA methylation levels than in wt EpiLCs. Percentage of total CpG DNA methylation (5mCpG) as measured by RRBS. Horizontal black lines within boxes represent median values, boxes indicate the upper and lower quartiles, and whiskers indicate the 1.5 interquartile rangeDashed red line indicates the median mCpG methylation in wt EpiLCs. **c**, Principal component (PC) analysis of RRBS data from wt, T1CM, T2CM, and T12CM ESCs and wt EpiLCs. **d**, PC analysis of RNA-seq data from wt, T1CM, T2CM, and T12CM ESCs during EpiLC differentiation.

**Supplementary Figure 3:**
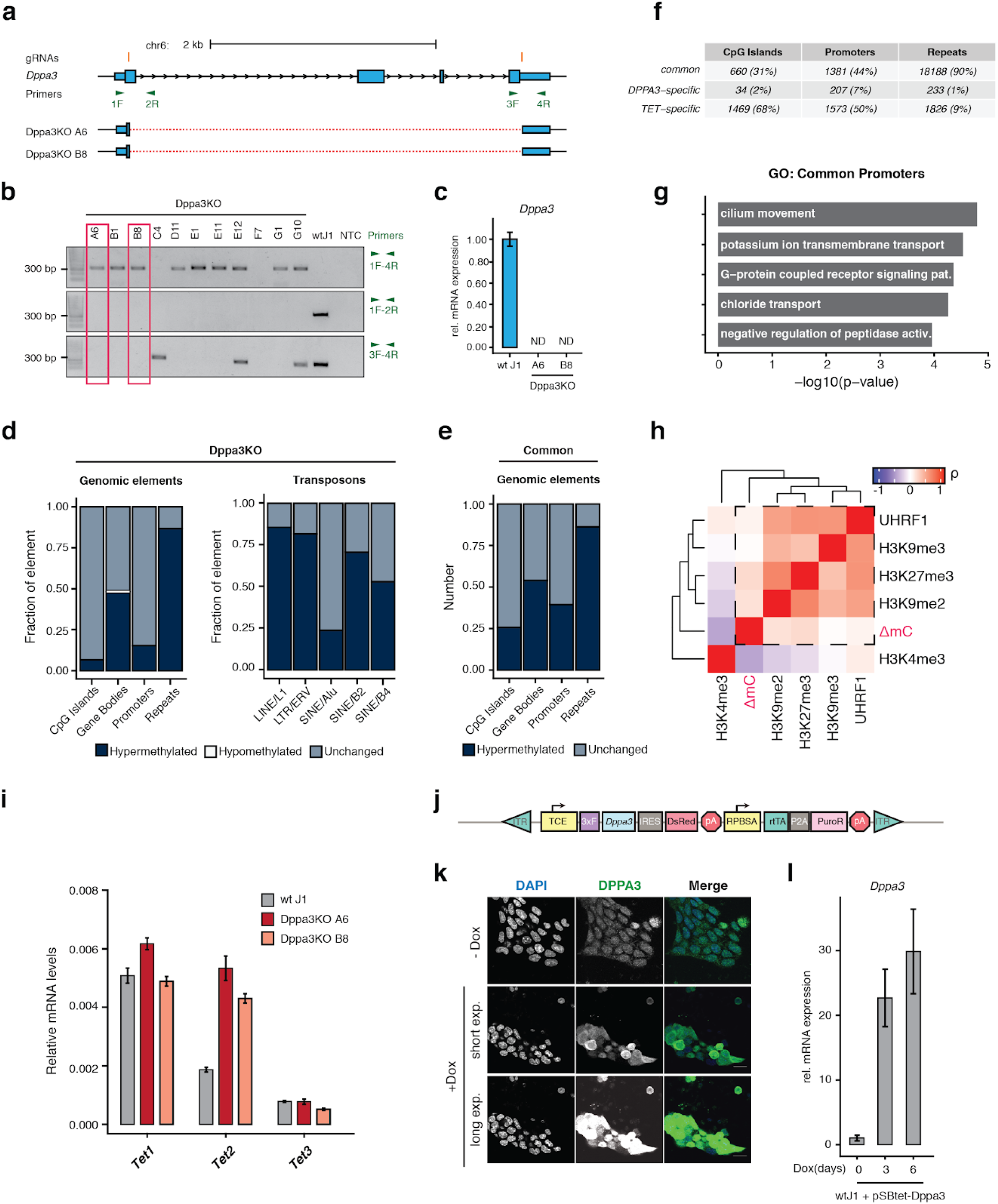
Generation, characterization, and methylome profiling of Dppa3KO ESCs. **a**, Schematic representation of the CRISPR/Cas9 gene editing strategy used to excise the entire *Dppa3* locus. The position of the two locus-flanking gRNAs are shown in orange. PCR primers for determination of locus removal and zygosity are indicated in green. **b**, Results of PCR using the primers indicated in (**a**). *Dppa3* knockout (Dppa3KO) clones, A6 and B8, chosen for further experiments are indicated by the red boxes. The wild-type (wt) and no template control (NTC) reactions are depicted on the right. The sequence alignment of the amplicon generated using primers 1 and 4 for Dppa3KO clones A6 and B8 is provided in the lower portion of (**a**) with solid boxes and the dashed lines representing successfully aligned sequences and genomic sequences not found in the sequenced amplicons respectively. **c**, *Dppa3* transcript levels of the two Dppa3KO clones A6 and B8 are depicted relative to mRNA levels in wt J1 ESCs after normalization to *Gapdh*. Error bars indicate mean ± SD calculated from technical triplicate reactions from *n* = 4 biological replicates. N.D., expression not detectable. **d**, Relative proportion of DNA hypermethylation occurring at each genomic element or retrotransposon class in Dppa3KO ESCs. **e**, Relative proportion of DNA hypermethylation common to T1CM, T2CM, and T12CM ESCs and Dppa3KO ESCs at each genomic element. **f**, Summary of differentially methylated regions either unique to TET mutants (TET-specific) and Dppa3KO ESCs (DPPA3-specific) or shared among TET mutants and Dppa3KO ESCs (common). **g**, Gene ontology (GO) terms associated with promoters specifically dependent on TET activity; adjusted p-values calculated using Fisher’s exact test. **h**, Cross-correlation analysis between DNA hypermethylation (ΔmC) in Dppa3KO ESCs and genomic occupancy of histone modifications and UHRF1 binding. Correlations are calculated over 1kb tiles. Positive correlations with ΔmC are bounded by a dashed-line square. **i**, Hypermethylation in Dppa3 KO ESCs is not associated with *Tet* downregulation. Expression of *Tet* genes in wt Dppa3KO ESC clones depicted as mRNA levels relative to *Gapdh*. Error bars indicate mean ± SD calculated from technical triplicate reactions from *n* = 2 biological replicates of each genotype or clone depicted. **j**, Schematic representation of the pSBtet-D3 (pSBtet-3xFLAG-Dppa3-IRES-DsRed) cassette for the Sleeping Beauty transposition-mediated generation of doxycycline (Dox) inducible *Dppa3* ESC lines. Abbreviations: inverted terminal repeat (ITR), tetracycline response element plus minimal CMV (TCE), 3xFLAG tag (3xF), internal ribosomal entry site (IRES), polyA signal (pA), constitutive RPBSA promoter (RPBSA), reverse tetracycline-controlled transactivator (rtTA), self-cleaving peptide P2A (P2A), puromycin resistance (PuroR). **k**, Confirmation of DPPA3 protein induction as assessed by immunofluorescence in uninduced ESCs (-Dox) or after 24 h of doxycycline treatment (+Dox). To illustrate the increase in DPPA3 protein levels, the same acquisition settings used to detect DPPA3, including a long exposure, in uninduced cells were applied for detection of DPPA3 after induction (+Dox (long exp.); bottom panel) leading to a saturated signal. Shorter exposure settings were also applied to the induced cells (+Dox (short exp.); middle panel) to better resolve the localization of DPPA3 after induction. The staining was repeated in 3 biological replicates with similar results. Scale bar: 10 µm. **l**, *Dppa3* expression before induction and after 3 or 6 days of doxycycline treatment in wt J1 + pSBtet-D3 ESCs. mRNA levels of *Dppa3* are shown relative to those in uninduced wt J1 ESCs after normalization to *Gapdh*. Error bars indicate mean ± SDcalculated from technical triplicate reactions from *n* = 2 biological replicates. In **d-h**, hypermethylation is a gain in 5mC compared to wt ESCs (q value < 0.05; absolute methylation difference > 20%).

**Supplementary Figure 4:**
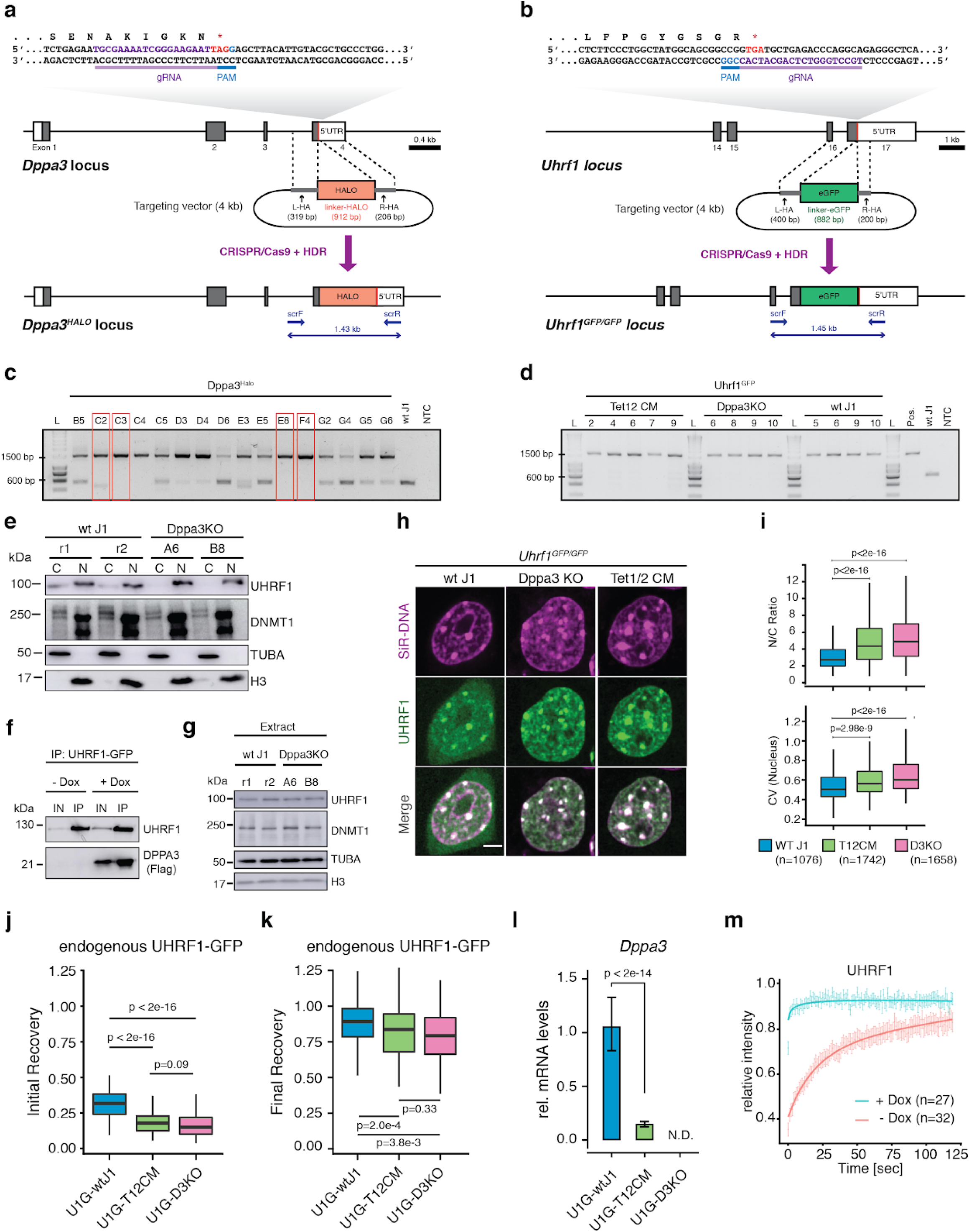
DPPA3 hinders UHRF1 chromatin association in mouse ESCs. **a,b**, Schematic of the CRISPR/Cas9 targeting strategy used to insert fluorescent protein tags at the (**a**) *Dppa3* and (**b**) *Uhrf1* loci. Exons are depicted as boxes, with the coding sequences shaded in grey. Stop codons are indicated with a red line. The nucleotide and amino acid sequence encompassing the gRNA target site are enlarged for detail. gRNAs target sequences are colored purple and their respective PAMs blue. Donor constructs harboring the coding sequence for (**a**) HALO or (**b**) eGFP are flanked by homology arms (L-HA, left homology arm and R-HA, right homology arm) and indicated as grey rectangles. Genotyping PCR primers used in (**c,d**) are shown as arrows along with the size of the amplicon confirming successful integration of the donor sequence. **c,d**, Results of genotyping PCRs for confirming the successful integration of (**c**) HALO into the *Dppa3* locus of wt J1 ESCs and (**d**) eGFP into the *Uhrf1* locus of wt J1, Dppa3KO, and T12CM ESCs. Wild-type (wt) and negative control without DNA template (NTC) are depicted on the right. **c**, Clones with Sanger-sequence validated homozygous insertion of the HALO coding sequence are indicated with red boxes. **d**, PCR results after pre-screening via microscopy. All depicted clones have a correct homozygous insertion of eGFP as validated by Sanger-sequencing. PCRs were performed using the primers depicted in (**a,b**). **e**, DPPA3 regulates subcellular UHRF1 distribution. Cytoplasmic, “C”, and nuclear, “N”, fractions from wt and Dppa3KO ESCs analyzed by immunoblot detection using the indicated antibodies. The fractionation and immunoblot were repeated 2 times with similar results obtained. **f**, Immunoblot detection of anti-GFP immunoprecipitated material (IP) and the corresponding input (IN) extracts from U1G/wt + pSBtet-D3 ESCs treated before (-Dox) and after (+Dox) 24 h induction of *Dppa3* with doxycycline. Immuoblot was repeated 2 times with similar results obtained. **g**, UHRF1 and DNMT1 protein levels are unaffected by DPPA3 loss. Whole-cell extracts from wt and Dppa3KO ESCs analyzed by immunoblot detection using the indicated antibodies. Immunoblot depicts *n* = 2 biological replicates. **h,i**, DPPA3 alters the localization and nuclear patterning of endogenous UHRF1-GFP. **h**, Representative confocal images of UHRF1-GFP in live U1G-wt, U1G-T12CM, U1G-Dppa3KO ESCs with DNA counterstain (SiR-Hoechst). Scale bar: 5 μm. **i**, Quantification of endogenous UHRF1-GFP (top panel) Nucleus to cytoplasm ratio (N/C Ratio) and (bottom panel) coefficient of variance (CV) within the nucleus of cells (number indicated in the plot) from *n* = 3 biological replicates per genotype. The assay was repeated 2 times with similar results. **j,k**, Endogenous DPPA3 prevents excessive UHRF1 chromatin binding in ESCs. Further analysis of the single cell FRAP data presented in Fig. 4c. showing the (**j**) initial and (**k**) final relative recovery of endogenous UHRF1-GFP. Intensity measurements 1 s and 60 s after photobleaching were used for calculating (**j**) initial recovery and (**k**) final recovery, respectively. **l**, *Dppa3* is significantly downregulated in U1G-T12CM ESCs. *Dppa3* transcript levels in U1G-wt, U1G-T12CM, U1G-Dppa3KO ESCs are depicted relative to mRNA levels in U1G-wt J1 ESCs after normalization to Gapdh. Error bars indicate mean ± SD calculated from technical triplicate reactions from *n* = 4 biological replicates (independent clones). N.D., expression not detectable. **m**, DPPA3 expression abolishes UHRF1 chromatin binding. FRAP analysis of endogenous UHRF1-GFP before (-Dox) and after 48 h of *Dppa3* induction (+Dox) in U1G-wt + pSBtet-D3 ESCs. The FRAP experiment was repeated at least 3 times with similar results obtained. In the boxplots in **i-k**, darker horizontal lines within boxes represent median values. The limits of the boxes indicate upper and lower quartiles, and whiskers indicate the 1.5-fold interquartile range. P-values in **i-l**; Welch’s two-sided t-test. In **m**, the mean fluorescence intensity of *n* cells (indicated in the plots) at each timepoint depicted as a shaded dot. Error bars indicate mean ± SEM. Curves (solid lines) indicate double-exponential functions fitted to the FRAP data.

**Supplementary Figure 5:**
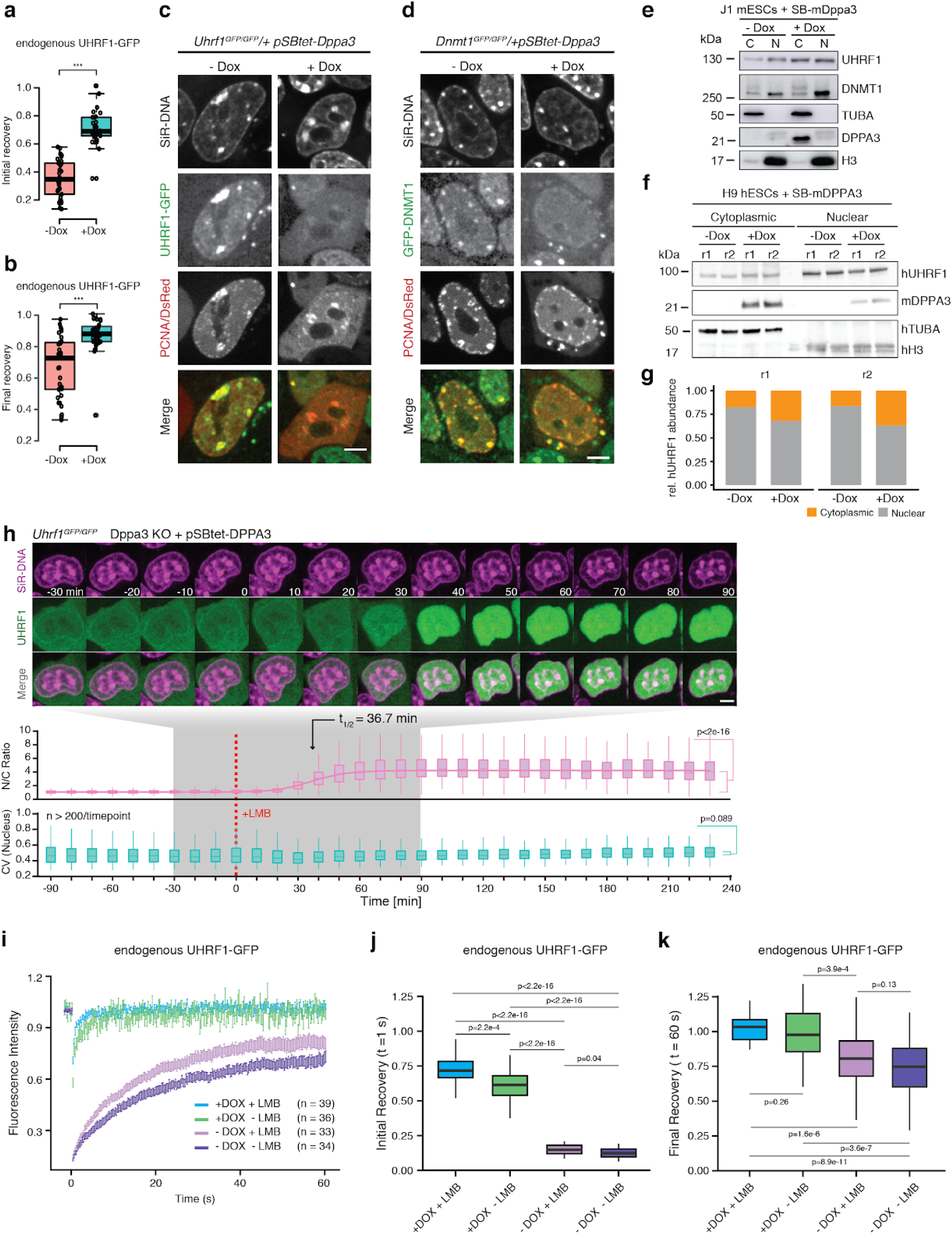
DPPA3 alters the localization and chromatin binding of endogenous UHRF1 and DNMT1 in ESCs. **a,b**, DPPA3 releases UHRF1 from chromatin. Initial (**a**) and final (**b**) relative recovery of UHRF1-GFP in *Uhrf1*^*GFP/GFP*^*/SBtet-3xFLAG-Dppa3* ESCs before (-Dox) and after 48 h of *Dppa3* induction (+Dox) (+Dox: *n* = 27; -Dox: *n* = 32) (data from FRAP experiment in Supplementary Fig. 4m). **c,d**, DPPA3 disrupts the recruitment of UHRF1 and DNMT1 to replication foci. Live cell imaging illustrating the late S-phase localization of (**c**) UHRF1-GFP in U1G/wt + pSBtet-D3 ESCs or (**d**) GFP-DNMT1 in live *Dnmt1*^*GFP/GFP*^ + pSBtet-D3 ESCs before (-Dox) and after 48 h of *Dppa3* induction. Transfected RFP-PCNA marks sites of active DNA replication within the nucleus. The assay was performed 2 times with similar results. Expressed free DsRed is a marker of doxycycline induction (see cytoplasmic signal in +Dox PCNA/DsRed panels and Supplementary Fig. 3j). DNA was stained in live cells using a 30 min treatment of SiR-DNA (SiR-Hoechst). Scale bar: 5 μm. **e-g**, DPPA3 alters the subcellular distribution of endogenous UHRF1 in mouse ESCs and human ESCs. **e,f**, Immunoblots of nuclear, “N”, and cytoplasmic, “C”, fractions from (**e**) U1G/wt + pSBtet-D3 mouse ESCs and (**f**) Human H9 ESCs + SBtet-3xFLAG-Dppa3 before (-Dox) and after 24 hours of *Dppa3* induction (+Dox) using the indicated antibodies. An anti-FLAG antibody was used for detection of FLAG-DPPA3. **g**, Quantification of the relative abundance of hUHRF1 in the cytoplasmic and nuclear fractions shown in (**f**) with the results of two independent biological replicates (r1 and r2) displayed. **h**, Nuclear export is dispensable for DPPA3-mediated inhibition of UHRF1 chromatin association. Localization dynamics of endogenous UHRF1-GFP in response to inhibition of nuclear export using leptomycin-B (LMB) after *Dppa3* induction in U1G/D3KO + pSBtet-D3 ESCs with confocal time-lapse imaging over 5.5 h (10 min intervals). *t = 0* corresponds to start of nuclear export inhibition (+LMB). (*top panel*) Representative images of UHRF1-GFP and DNA (SiR-Hoechst stain) throughout confocal time-lapse imaging. Scale bar: 5 μm. (middle panel) Nucleus to cytoplasm ratio (N/C Ratio) of endogenous UHRF1-GFP signal. (bottom panel) Coefficient of variance (CV) of endogenous UHRF1-GFP intensity in the nucleus. (*middle and bottom panel*) N/C Ratio and CV values: measurements in *n* > 200 single cells per time point, acquired at *n* = 20 separate positions. Curves represent fits of four parameter logistic (4PL) functions to the N/C Ratio (pink line) and CV (green line) data. The experiment was repeated 2 times with similar results. **i-k**, Nuclear export is not only dispensable for but attenuates DPPA3-mediated inhibition of UHRF1-GFP chromatin binding. **i**, FRAP analysis of endogenous UHRF1-GFP within the nucleus of U1G/D3KO + pSBtet-D3 ESCs before (-Dox) and after 48 h of *Dppa3* induction and before (-LMB) and after (+LMB) 3 h of leptomycin-B (LMB) treatment. The mean fluorescence intensity of *n* cells (indicated in the plots) at each timepoint depicted as a shaded dot. Error bars indicate mean ± SEM. The FRAP experiment was repeated 3 times with similar results. **j,k**, Further analysis of the data from (**i**) showing the (**j**) initial and (**k**) final relative recovery of endogenous UHRF1-GFP. For **a,b,j,k**, initial and final recoveries were calculated using the intensities measured 1 s or 60 s after photobleaching, respectively. In the boxplots, darker horizontal lines within boxes represent median values. The limits of the boxes indicate upper and lower quartiles, and whiskers indicate the 1.5-fold interquartile range. Welch’s two-sided t-test. *** P < 0.001; ** P = 0.002; * P = 0.03; n.s.: not significant.

**Supplementary Figure 6:**
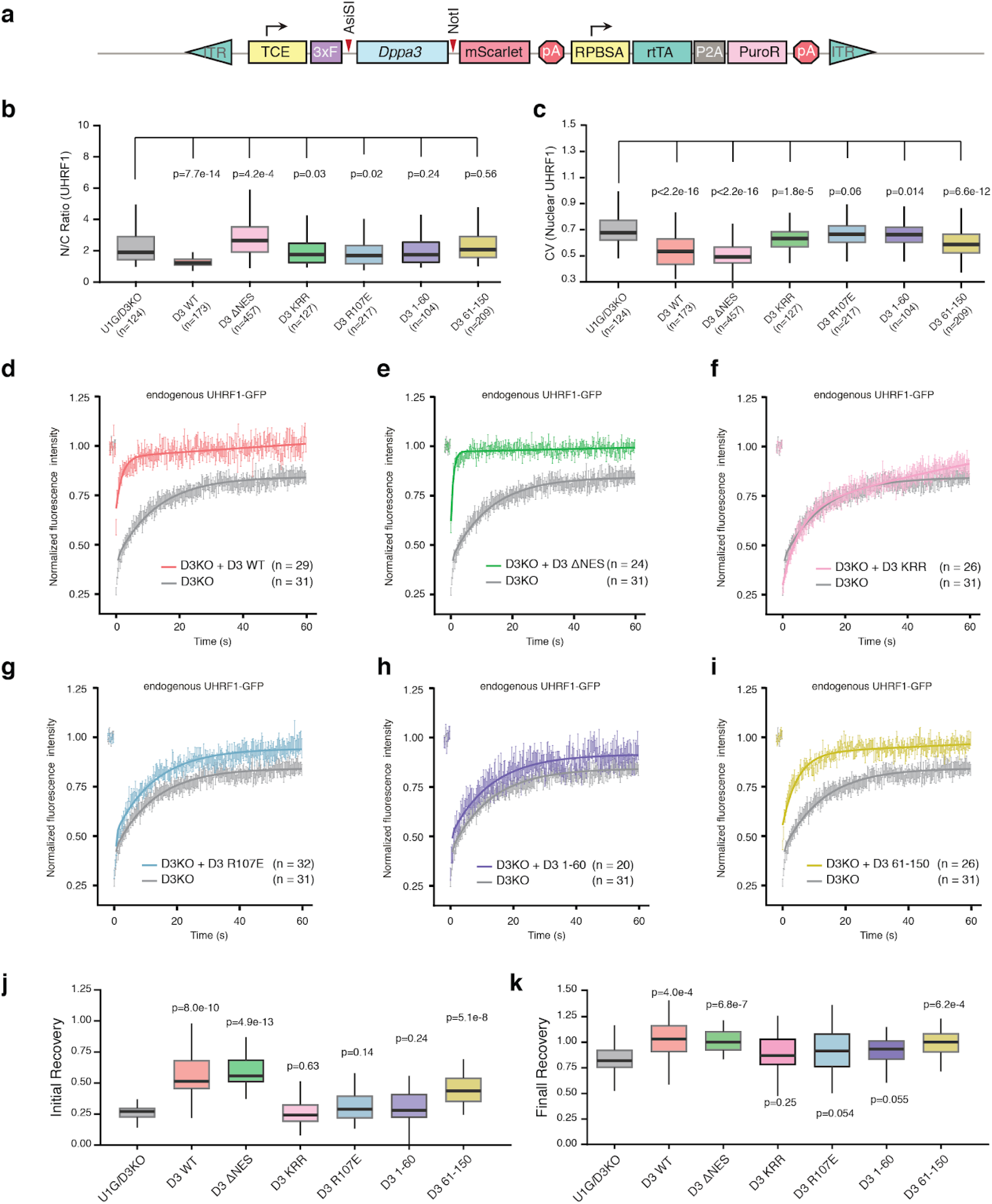
Further analysis of endogenous UHRF1-GFP localization and chromatin binding in response to mutant forms of mDPPA3. **a**, Schematic representation of the pSBtet-D3-mSC (pSBtet-3xFLAG-Dppa3-mScarlet) cassette for the Sleeping Beauty transposition-mediated generation of ESC lines with doxycycline (Dox) inducible expression of DPPA3-mScarlet fusions. Abbreviations: inverted terminal repeat (ITR), tetracycline response element plus minimal CMV (TCE), 3xFLAG tag (3xF), internal ribosomal entry site (IRES), polyA signal (pA), constitutive RPBSA promoter (RPBSA), reverse tetracycline-controlled transactivator (rtTA), self-cleaving peptide P2A (P2A), puromycin resistance (PuroR). Restriction sites used for exchanging the *Dppa3* coding sequences are indicated with red arrows. **b,c**, DPPA3 alters the nuclear localization of UHRF1 independently of promoting UHRF1 nucleocytoplasmic translocation. Quantification of endogenous UHRF1-GFP (**b**) Nucleus to cytoplasm ratio (N/C Ratio) and (**c**) coefficient of variance (CV) within the nucleus of U1G/D3KO + pSBtet-D3 ESCs expressing the indicated mutant forms of *Dppa3. n* = number of cells analyzed (indicated in the plot). **d-i** Nuclear export and the N-terminus of DPPA3 are dispensable for its inhibition of UHRF1 chromatin binding. FRAP analysis of endogenous UHRF1-GFP within the nucleus of U1G/D3KO + pSBtet-D3 ESCs expressing the following forms of DPPA3: (**d**) wild-type (D3 WT), (**e**) nuclear export mutant L44A/L46A (D3-ΔNES), (**f**) K85E/R85E/K87E mutant (D3-KRR), (**g**) R107E mutant (D3-R107E), (**h**) N-terminal 1-60 fragment (D3 1-60), (**i**) C-terminal 61-150 fragment (D3 61-150). In **d-i**, the mean fluorescence intensity of *n* cells (indicated in the plots) at each timepoint is depicted as a shaded dot. Error bars indicate mean ± SEM. Curves (solid lines) indicate double-exponential functions fitted to the FRAP data. Individual same fits correspond to those in Fig. 5c. FRAP of UHRF1 in cells expressing *Dppa3* mutants was repeated 2 times. **j,k**, Further analysis of the data from (**d-i**) showing the (**j**) initial and (**k**) final relative recovery of endogenous UHRF1-GFP. Intensity measurements 1 s and 60 s after photobleaching were used for calculating (**j**) initial recovery and (**k**) final recovery, respectively. In the boxplots in **b, c, j**, and **k**, darker horizontal lines within boxes represent median values. The limits of the boxes indicate upper and lower quartiles, and whiskers indicate the 1.5-fold interquartile range. P-values based on Welch’s two-sided t-test of the difference between U1/D3KO and each DPPA3 mutant.

**Supplementary Fig. 7:**
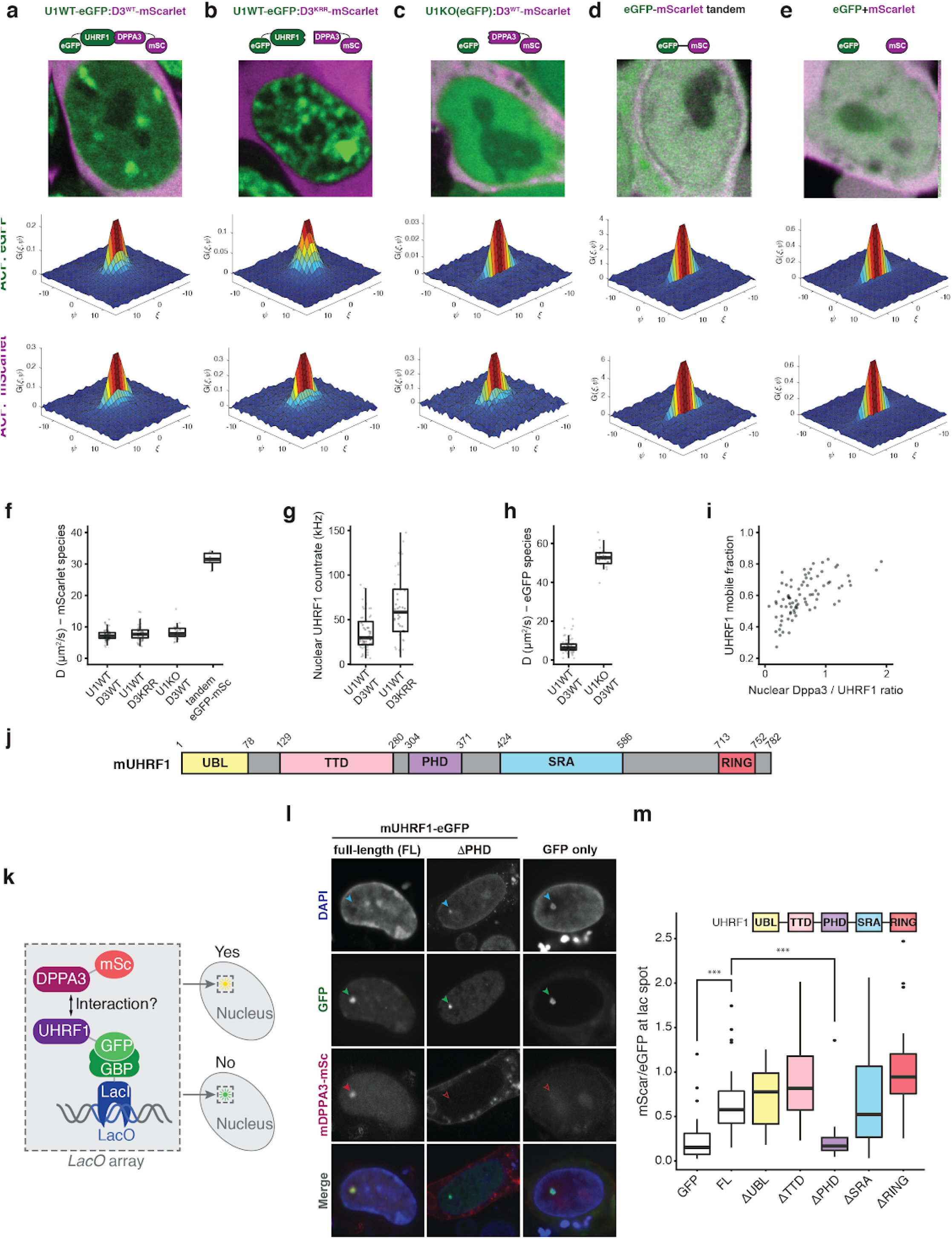
DPPA3 binds UHRF1 via its PHD domain to form a mobile complex in ESCs. **a-e**, Representative dual-color live confocal images (top row) and associated 3D plots of mean autocorrelation (ACF) values of eGFP (middle row) and mScarlet (bottom row) species in U1G/D3KO + pSBtet-D3 ESCs expressing the following forms of DPPA3-mScalet: (**a**) wild-type (U1WT:D3^WT^) and (**b**) K85E/R85E/K87E mutant (U1WT:D3^KRR^), in (**c**) Uhrf1KO ESCs expressing free eGFP and wild-type *Dppa3*-mScarlet (U1KO:D3^WT^), and in control ESCs expressing (**d**) an eGFP-mScarlet tandem fusion (eGFP-mScarlet) and (**e**) free eGFP and free mScarlet (eGFP + mScarlet). In **a-e**, each image is the result of merging the sum projections of 250-frame image series acquired simultaneously from eGFP (green) and mScarlet (magenta) channels. Autocorrelation plots are color-coded to indicate mean correlation value. The fast timescale axis is indicated by *ξ*, and the slow timescale axis is indicated by *ψ*. **f**, Calculated diffusion coefficient of mScarlet species in different cell types (those shown in (**a-d**)), derived from a two-component fit of mScarlet autocorrelation functions. Data are pooled from three independent experiments, except tandem eGFP-mScarlet which was from two independent experiments. **g**, Photon count rates of UHRF1-eGFP in nuclei of U1G/D3KOs expressing either DPPA3-WT (U1WT:D3^WT^) or DPPA3-KRR mutant (U1WT:D3^KRR^). Data are pooled from three independent experiments. **h**, Calculated diffusion coefficients of UHRF1-GFP (in U1WT:D3^WT^) or free eGFP (in U1KO:D3^WT^), derived as those in C. Data are pooled from three independent experiments **i**, Scatter plot showing the relationship between the mobile fraction of UHRF1-eGFP and the ratio of DPPA3-mScarlet:UHRF1-eGFP photon count rates in the nucleus. Data are pooled from three independent experiments. **j**, A schematic diagram illustrating the domains of mUHRF1: ubiquitin-like (UBL), tandem tudor (TTD), plant homeodomain (PHD), SET-and-RING-associated (SRA), and really interesting new gene (RING). **k**, Overview of the F3H assay used to find the domain of UHRF1 which binds to DPPA3. **l,m**, The PHD domain of UHRF1 is necessary for mediating the interaction with DPPA3. **l**, Representative confocal images of free GFP or the ΔPHD UHRF1-GFP constructs (**j**) immobilized at the lacO array (indicated with green arrows). Efficient or failed recruitment of DPPA3-mScarlet to the lacO spot are indicated by solid or unfilled red arrows respectively. **m**, The efficiency of DPPA3-mScarlet recruitment to different UHRF1-eGFP deletion constructs immobilized at the lacO array is given as the fluorescence intensity ratio of mScarlet (DPPA3) to eGFP (UHRF1 constructs) at the nuclear LacO spot. Data represents the results of *n* > 10 cells analyzed from two independent experiments. In **f-h**, each data point indicates the measured and fit values from a single cell. In the box plots **f-h** and **m**, darker horizontal lines within boxes represent median values. The limits of the boxes indicate upper and lower quartiles, and whiskers indicate the 1.5-fold interquartile range. In **m**, P-values were calculated using Welch’s two-sided t-test: ** P < 0.01; *** P < 0.001.

**Supplementary Figure 8:**
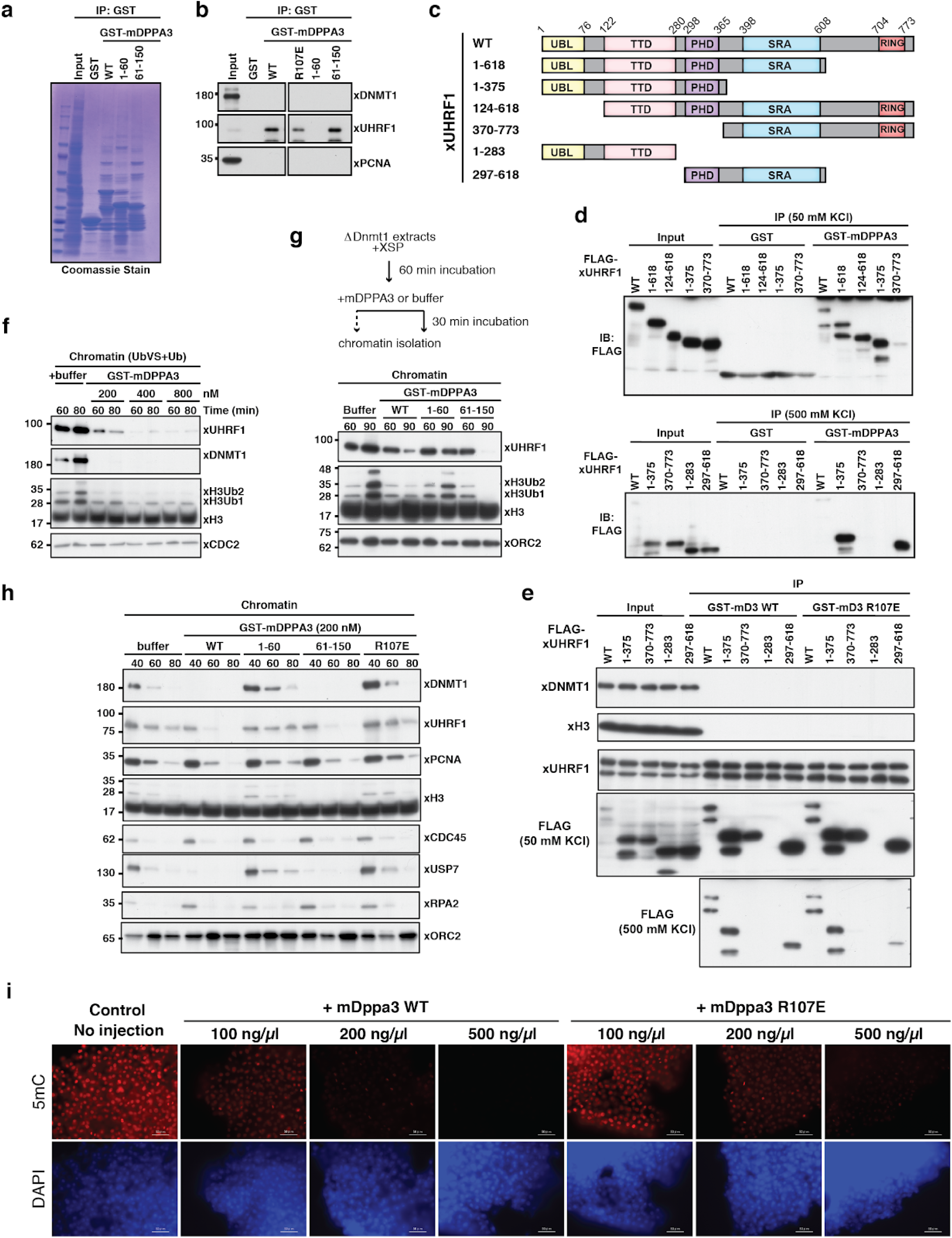
DPPA3 evolved in boreoeutherian mammals but its function is conserved in lower vertebrates. **a**, mDPPA3 C-terminus is sufficient for xUHRF1 binding. GST-tagged mDPPA3 wild-type (WT) and truncations (1-60 and 61-150) were immobilized on GSH beads and incubated with *Xenopus* egg extracts. The samples shown in Fig. 7c were subjected to SDS-PAGE and stained with Coomassie Blue. **b**, High affinity interaction between mDPPA3 and xUHRF1 is weakened by R107E. GST-tagged mDPPA3 wild-type (WT), the point mutant R107E, and truncations (1-60 and 61-150) were immobilized on GSH beads and incubated with *Xenopus* egg extracts. Pull-downs were subjected to stringent 500 mM KCl washing. Bound proteins were analyzed using the indicated antibodies. **c**, Schematic diagram illustrating the xUHRF1 deletion constructs used in the pull-downs in (**d,e**). **d,e**, xUHRF1 binds mDPPA3 via its PHD domain. *In vitro* translated xUHRF1 fragments were added to interphase *Xenopus* egg extracts. (**d**) GST and GST-mDPPA3 wild-type (WT) or (**e**) GST-tagged mDPPA3 wild-type (WT) and GST-tagged mDPPA3 R107E were immobilized on GSH beads and then used for GST-pulldowns on egg extracts containing the indicated recombinant xUHRF1 fragments. Bound proteins were analyzed using the denoted antibodies. Pull-downs were subjected to either 50 mM KCl or more stringent 500 mM KCl washing as indicated. **f**, mDPPA3 disrupts xUHRF1-dependent ubiquitylation of H3. As dual-monoubiquitylation of H3 (H3Ub2) is hard to detect given its quick turnover^89^, we specifically enhanced ubiquitylation by simultaneous treatment of extracts with ubiquitin vinyl sulfone (UbVS), a pan-deubiquitylation enzyme inhibitor^149^ and free ubiquitin (+Ub) as described previously^89^. Buffer or the displayed concentration of recombinant mDPPA3 were then added to the extracts. After the indicated times of incubation, chromatin fractions were isolated and subjected to immunoblotting using the antibodies indicated. **g**, mDPPA3 displaces chromatin-bound xUHRF1. To stimulate xUHRF1 accumulation on hemi-methylated chromatin, *Xenopus* extracts were first immuno-depleted of xDNMT1 as described previously^66^. After addition of sperm chromatin, extracts were incubated for 60 min to allow the accumulation of xUHRF1 on chromatin during S-phase and then the indicated form of recombinant mDPPA3 (or buffer) was added. Chromatin fractions were isolated either immediately (60 min) or after an additional 30 min incubation (90 min) and subjected to immunoblotting using the antibodies indicated. **h**, The region 61-150 of mDPPA3 is sufficient but requires R107 to inhibit xUHRF1 chromatin binding. Sperm chromatin was incubated with interphase *Xenopus* egg extracts supplemented with buffer (+buffer) and GST-mDPPA3 wild-type (WT), the point mutants R107E, truncations (indicated). Chromatin fractions were isolated after the indicated incubation time and subjected to immunoblotting using the antibodies indicated. **i**, mDPPA3-mediated DNA demethylation in medaka. Representative 5mC immunostainings in late blastula stage (∼8 h after fertilization) medaka embryos injected with wild-type *mDppa3* (WT) or *mDppa3* R107E (R107E) at three different concentrations (100 ng/µl, 200 ng/µl, or 500 ng/µl). Scale bars represent 50 µm. DNA counterstain: DAPI,4′,6-diamidino-2-phenylindole.

